# Urban effects on timing and variability of diel activity differ across passerine species and seasons

**DOI:** 10.1101/2024.12.19.629353

**Authors:** Pablo Capilla-Lasheras, Claire J. Branston, Paul Baker, Cara Cochrane, Barbara Helm, Davide M. Dominoni

## Abstract

1. Life on Earth is adapted to rhythmic cycles in environmental conditions throughout the day and year via diel patterns of behavioural activity.

2. Urban conditions can disrupt such behavioural rhythms of activity. However, most studies so far have investigated urban effects on patterns of activity of single species in a single season. Additionally, we know little about the level of between and within-individual variation in urban and non-urban populations, and whether they differ.

3. Here, we use automated radio telemetry to record patterns of daily activity in six passerine species (blackbird, robin, great tit, blue tit, dunnock and chaffinch) across two urban and two forest populations during the pre-breeding and post-breeding season. We investigate urban effects on five activity-related traits: time of activity onset, time of activity end, duration of diurnal activity, level of diurnal activity and level of nocturnal activity. We employ statistical tools that allow us to estimate urban effects on mean phenotypic values, but also quantify urban *versus* forest differences in between-individual and within-individual phenotypic variation.

4. We found the strongest urban effects on time of activity onset in blackbirds and robins during the pre-breeding season: urban populations of blackbird and robin started their daily activity earlier than their forest counterparts. We did not find this effect in the other species. Urban populations of all species showed higher levels of nocturnal activity than forest populations, but this effect was not offset by lower diurnal activity, suggesting that urban birds may incur higher daily energetic demands.

5. Lastly, our analysis revealed large and consistent differences in variation in the investigated timing traits. Urban birds were more variable between individuals, implying lower population synchronization, and more variable within individuals, implying less consistent behaviour than their forest counterparts. Conversely, activity levels were more variable in forest birds.

6. We conclude that, for birds, urban life is associated with less rest, less consistency, and lower synchronicity, but that effect sizes depend on species and time of the year. Our results warn against generalising the effects of urbanisation on daily rhythms of birds, and call for future studies to understand the mechanisms behind species and seasonal differences.

## Introduction

As the Earth rotates, environmental conditions cycle through predictable daily phases. Species have adapted to these predictable fluctuations in environments and, indeed, diel rhythms of activity are nearly universal (Foster & Kreitzman, 2005). Appropriate timing of activity is crucial for survival and optimal reproduction, and several mechanisms allow organisms to track diel changes in environments, thereby fine-tuning their behaviour to optimally match environmental conditions (Helm et al., 2017). Factors associated with urbanisation (e.g., artificial light at night, noise pollution, perceived predation risk, or changes in food webs) frequently mimic the natural cues that organisms use to time their activity (Dominoni et al., 2020), changing temporal environments and leading to changes in behavioural rhythms of organisms living in urban habitats (e.g., Gaynor et al., 2018). Despite evidence for differences in behavioural timing of urban populations, we still have a poor understanding of their eco-evolutionary significance. Behavioural differences in how animals time their daily activity can have ecological impacts (e.g., promote new interactions between species; (Kronfeld-Schor & Dayan, 2003; Martinez-Bakker & Helm, 2015) and evolutionary implications, as rhythmic behavioural traits are correlated with fitness proxies in the wild (e.g., Kempenaers et al., 2010; Womack et al., 2023). We lack comparative studies that investigate multiple species throughout the year, assess between and within-individual variation in patterns of activity, and examine how these sources of variation are shaped by biological and environmental factors.

Changes in behavioural rhythms associated with urban living have been shown in several species and can be large (Gilbert et al., 2023). In mammals, for example, human activity is positively related to the degree of nocturnality (Gaynor et al., 2018). Similarly, multiple lines of evidence suggest substantial changes in behavioural and physiological rhythms of urban birds. Urban birds start their daily activity earlier than non-urban populations (Dominoni et al., 2013; McGlade et al., 2023; Silva et al., 2015). In many cases, altered rhythms have been linked to artificial light at night (ALAN; Sanders et al., 2020; Sun et al., 2017), but other factors, such as disturbance and noise, have also been shown to affect behavioural rhythms (Connelly et al., 2020; De Framond & Brumm, 2022; Gaynor et al., 2018; Gilbert et al., 2023). Experimental data confirm correlative observations and provide evidence for parallel physiological changes (Dominoni et al., 2013, 2022). Birds are a key focus of research on how urban conditions affect biological rhythms because they form rich communities with species of variable biological features. Most avian species are diurnal, like humans (unlike most urban mammals), and they can carry biologging tools to record their activity. In diurnal avian species, it is well documented that urban conditions are associated with the extension of daytime activity into the night, occurring mostly in the morning; urban effects on activity timing in the evening are more limited (McGlade et al., 2023; Womack et al., 2023). Despite these well-documented patterns, several knowledge gaps still exist, hindering our ability to predict and fully understand how urban conditions impact the rhythms of activity of wild animals.

First, previous studies mostly focus on single species, but sensitivity of rhythmic behavioural traits to urban conditions might differ between species. Indeed, studies on timing of singing across passerine species in Europe have reported that early morning singing species (e.g., robins [*Erithacus rubecula*] or blackbirds [*Turdus merula*]) advance their dawn song in ALAN-polluted areas, whereas later singing species (e.g., chaffinch [*Fringilla coelebs*]) do not show this response (Kempenaers et al., 2010; Silva et al., 2014). However, these studies used dawn and dusk singing as measurements of behavioural rhythms, but singing is only a limited facet of daily activities and at temperate latitudes, largely restricted to males. Conversely, recent studies have measured activity rhythms during incubation of female birds and also reported advances under ALAN, but no comparable data exist for males (McGlade et al., 2023; Womack et al., 2023). In two species of tits (great tit [*Parus major*] and blue tit [*Cyanistes caeruleus*]), experimental application of ALAN altered roosting behaviour, with species-specific response patterns (Sun et al., 2017). Thus, there is indication of species-specific and sex-specific responses of birds to urban-linked environmental conditions (including ALAN), but data are patchy and currently difficult to interpret.

Second, previous studies looking at the effects of urban conditions on rhythms of activity have been mostly conducted during or just before the breeding season (but see Raap et al., 2015). The breeding season represents only a portion of the annual cycle, when both intrinsic (e.g., physiological state) and extrinsic conditions are profoundly different from those in other periods of the year. In temperate areas, winter nights are long and deciduous trees are leafless, which may increase exposure to ALAN. Investigating urban effects across contrasting periods of the year could yield a more complete picture. Physiological state might drive seasonal differences in the response of daily rhythms to the urban environment. A season-specific advance of activity was also suggested by avian captivity studies, which indicated earlier timing of activity in spring compared to autumn (Daan & Aschoff, 1975). Experimental studies also demonstrated direct effects of testosterone on avian biological rhythms (Gwinner, 1975). In males, the strong morning advancement in locomotor and singing activity coincides with the courtship and mating phase, when levels of circulating androgens are high and when early dawn song may be rewarded by higher extra-pair paternity (Poesel et al., 2006). Thus, previous studies hypothesised that the urban environment might create an opportunity to advance dawn song earlier in the morning (e.g., via the presence of ALAN), and those individuals who can exploit the opportunity obtain fitness advantages. Such opportunities would be specific to the breeding season, and mechanisms such as effects of testosterone are heavily biased towards males (Gwinner, 1975), which also reinforces the need to quantify sex differences in activity timing. If males can gain reproductive advantages by advancing morning activity, we would expect stronger activity responses to urban conditions in males than in females (i.e., male advancements could be driven by intra-sexual competition; Kacelnik & Krebs, 1983).

Third, the major focus of research to date has been on mean patterns of activity timing, while comparing variation and consistency between and within urban and non-urban individuals has been largely neglected. Recent studies have suggested that urban conditions could lead to increases in phenotypic variation in a wide range of traits (CapillalJLasheras et al., 2022; Thompson et al., 2022). This is possibly due to increased micro-environmental heterogeneity in the urban environment (Capilla-Lasheras et al., 2022; Morelli et al., 2023), which could be partly created by patchy patterns of ALAN in these habitats. A previous study suggested that variation in activity timing was higher in urban than non-urban blackbirds, both within and between individuals, suggesting lower consistency and lower synchronicity of diel rhythms in the urban population (Dominoni et al., 2014).

Fourth, advancement of activity into the night and nocturnal restlessness have been reported to come at a cost of reduced night rest in experimental studies in birds that were experimentally exposed to ALAN or noise (Aulsebrook et al., 2020; Connelly et al., 2020; Dominoni et al., 2022). Sleep loss could contribute to the often-observed poorer condition of urban birds (Capilla-Lasheras et al., 2017; van Hasselt et al., 2021; Ouyang et al., 2017). However, for wild birds, individual-based data are still lacking for resolving whether 24 h activity levels differ between urban and non-urban birds because nocturnal sleep loss could be offset by day-time rest phases.

Here we use automated radio telemetry in two urban and two non-urban bird communities to holistically investigate how urban conditions impact patterns of activity across six species of songbirds. Using a large-scale telemetry deployment allowed us to acquire continuous measurements of activity around the clock for six species (European robin, European blackbird, great tit, blue tit, dunnock [*Prunella modularis*], and common chaffinch) in two distinct periods, pre-breeding (February-April) and post-breeding season (September-November), over two years. These six species were chosen to include natural variation in activity patterns, for example, in song timing (Silva et al., 2014), with early (e.g., robin and blackbirds) and late rising species (e.g., dunnock and chaffinch). In three species, we were also able to explore differences between the sexes. We investigated a) whether the effect of urbanisation on diel activity depended on the species; b) whether whether the effect of urbanisation on diel activity depended on sex; c) the effect of urbanisation on diel activity depended on the time of the year; d) whether urbanisation affected within and between-individual variation in diel activity; and, if urban birds showed higher levels of nocturnal activity, they compensated by resting more during the day.

Due to their natural predisposition to be active early, we predict that early rising species such as robins and blackbirds will show a stronger response (i.e., earlier time of onset of activity) to urbanisation than late rising species. Differences in the time of end of activity should be weaker or absent, as in previous studies (A). Due to intra-sexual competition, we would predict stronger effects of urban conditions on morning activity timing in males than in females (B). If extending activity in the morning is linked to reproductive physiology or fitness advantages, we predict stronger urban effects on activity patterns (particularly in the morning) during the pre-breeding than post-breeding phase (C). Based on previous findings of increased variation of the urban environment, we predict that the overall urban effects on mean timing traits will be accompanied by increases in both between and within-individual variation in time-keeping traits (D). Finally, due to direct human disturbance or effects of urban factors on sleep, we predict higher levels of nocturnal activity in urban birds. If increased nocturnal activity in urban habitats leads to increases in energy expenditure, we would expect reduced (i.e., compensatory) levels of diurnal activity (E).

## Methods

### Ethical note

All birds were caught, ringed and radio tagged after animal ethics clearance granted by the University of Glasgow and Nature Scot. Ringing and special methods licenses were granted by the British Trust for Ornithology to CJB and DMD. Institutional permission was granted in spring 2020 to continue work during periods of COVID-19 lockdown.

### Field methods: recording of activity of free-living birds

To estimate timing of activity and levels of activity in free-living birds, we use radio telemetry technology (Lotek, UK). Using mist-nets, we caught, individually marked with a metal ring (British Trust for Ornithology) and radio tagged (details below) individuals of six passerine species (european robin, eurasian blackbird, great tit, blue tit, dunnock, common chaffinch), during February-April (hereafter ‘pre-breeding phase’; range across both years = February 2 to April 16; mean = March 10; median = March 8) and September-November (hereafter ‘post-breeding phase’; range across both years = September 30 to November 5; mean = October 17; median = October 17) in 2020 and 2021 in two urban and two non-urban locations (Supplementary table 1). The length of each sampling period was determined by the time the radio receivers kept collecting data (details below). The urban sites were located within the boundaries of Glasgow city (Kelvingrove Park [55° 52.18N, 4° 17.22W] and Garscube Campus [55° 54.22N, 4° 19.2W]) and are dominated by non-native tree species, high levels of impervious surface and nearly constant human presence. The non-urban sites were located in the surroundings of the Scottish Centre for Ecology and the Environment (‘SCENE’), approximately 45 kilometres north of Glasgow, on the eastern shore of Loch Lomond (SCENE [56° 7.73N, 4° 36.79W] and Sallochy forest [56° 7.42N, 4° 36.07W]). Both non-urban sites are forests, dominated by native tree species. Passerine populations at these sites have been intensively studied in recent years and more detailed descriptions of these sites can be found in Branston et al., (2021), Jarrett et al., (2020) and Pollock et al., (2017).

Individuals of the six target species were radio tagged using Lotek Pip tags (Ag190 [for blue tits], Ag337 [for chaffinch, dunnock, great tit and robin] and Ag386 [for blackbird]; weight between 0.25 g [Ag190] and 2.20 g [Ag386]; hereafter referred to ‘tags’) mounted on the back of the birds using non-toxic glue and a piece of cotton fabric. Tags weighed in all cases less than 5% of the bird’s body weight. Tags continuously emitted radio pulses in a unique radio frequency allowing individuals to be followed and identified. In each site, we installed two to four automatic Lotek radio receivers (SRX800-D2) which were set to scan each tag (i.e., a unique frequency) every 180 seconds (three minutes). Each radio receiver was equipped with two to three uni-directional Yagi antennas, pointing in opposite directions and installed on rooftops or on dedicated 8-10 m masts. Receivers were installed at a mean distance of 239.00 metres (standard deviation = 196.63 meters; minimum distance between receivers = 65 metres; maximum distance between receivers = 697 meters). Whenever a tag was detected, the receiver recorded a date and time stamp, and the radio signal strength (dB). Birds were often out of range of receivers, and conversely, if present they were often recorded by more than one receiver. Similar approaches have been successfully employed to monitor patterns of activity in other bird species (Dominoni et al., 2013; Ouyang et al., 2017; Serota & Williams, 2019; Steiger et al., 2013).

### Radio telemetry data processing

The radio telemetry system generated a 3-min interval time series of radio detections per bird (i.e., tag) and per receiver. Following published studies using similar technology (e.g., Dominoni et al., 2014), we calculated the absolute difference in radio signal strength between consecutive time intervals (hereafter termed ‘signal differential’) per receiver and per bird. We then averaged this value across the receivers per site. This process produced a 3-min time series of signal differentials per bird that we used to infer daily onset of activity, end of activity and activity levels (Figure S1). The distribution of signal differentials was very similar per habitat and species, suggesting consistent power to estimate time of onset and end of activity (see below; Figure S2). Briefly, when a radio-tagged bird is static and its position in reference to receivers does not change (e.g., while roosting at night), we expect to see constant radio signal strength (i.e., signal differential close to zero); however, as soon as a radio-tagged bird becomes active, variation in its signal differential is expected to increase sharply (Figure S1).

### Estimation of timing and levels of activity

#### Estimation of daily onset and end of activity

We used frequentist change point analysis to estimate onset and end of activity based on the 3-min time series of signal differentials per bird (e.g., Dominoni et al., 2013). This method seeks to identify the point of change between the active and inactive state (see Supplementary methods for details). In short, for a given time series, this method splits the whole time-series in two blocks and a normal distribution is fitted to each data block. Then, the log maximum likelihood of the data is calculated and summed across both data blocks. The procedure is repeated moving the point for data split one time unit (i.e., 3 min) forward. This method results in a time series of log maximum likelihoods indicating the likelihood of a break in the distribution from which the data is originated; thereby providing statistical support for the location of the break in the time series between inactivity and activity. Based on previous research on daily patterns of activity in diurnal passerine birds (Dominoni et al., 2013; Maury et al., 2020; Womack et al., 2023), we restricted the estimation of daily onset of activity to a time window of six hours, between four hours before sunrise and two hours after sunrise. Similarly, for the estimation of end of activity, we restricted the analysis to an 8-hour time window between four hours before and after sunset. These time windows were chosen based on *a priori* knowledge of time of onset of activity in birds that tends to occur more often before sunrise than after sunrise, while time of end of activity is more varied and tends to occur more symmetrically around sunset (Dominoni et al., 2013, 2014). The choice of these windows was not dependent on habitat type or any other variable of interest. To control for temporal variation in sunrise and sunset times (which will strongly determine patterns of bird activity), we defined relative onset/end of daily activity as a bird’s onset/end of activity minus sunrise/sunset time (but see Figure S3 for a visualisation of absolute times of onset and end of activity over the recording periods). Therefore, positive values of relative onset/end of daily activity indicate that a bird started/ended its activity after sunrise/sunset. Negative values of relative onset/end of daily activity indicate that a bird started/ended its activity before sunrise/sunset. Duration of diurnal activity was then defined as the time difference between onset and end of activity per individual per day. Estimates of onset and end of activity were retained for downstream analysis when they came from morning/evening windows of analysis within which a given bird had been detected more than 75% of the total time in the window of interest (this produced a dataset for downstream analysis of 3,855 observations out of 5,679 raw data observations). Individuals of each species were monitored for a similar duration (blackbird = 14.6 [standard deviation, SD = 9.29] days; blue tit = 15.9 [SD = 10.10] days; chaffinch = 9.76 [SD = 9.07] days; dunnock = 13.2 [SD = 8.47] days; great tit = 13.2 [SD = 9.28] days; robin = 14.1 [SD = 8.29] days). Sample sizes (number of observations and individuals) included in the analysis per sex, species, year and habitat are provided in Table S1. We validated estimates of onset and end of activity via a second, independent, statistical method, Bayesian broken stick models (Supplementary methods 1 and Supplementary results 1). Both approaches led to similar conclusions. In the main text, we present results after applying the frequentist change point approach outlined above to estimate onset and end of activity. However, full results of both methods and their comparison can be found in Supplementary results 1, Figure S11, Table S2, Table S3.

#### Estimation of activity levels throughout the day

We classified each time point as ‘active’ or ‘inactive’ based on signal differentials. We assigned observations to ‘active’ when the difference in radio signal strength (i.e., signal differential) between consecutive time points was higher than 10 dB (Ouyang et al., 2017); similarly, we assigned observations to ‘inactive’ when the difference in radio signal strength between consecutive time points was lower than 10 dB (Ouyang et al., 2017). This signal differential threshold has been applied to great tits before (Ouyang et al., 2017) and while appropriate threshold values may differ between species, previous field validations suggest that the same threshold works well for similar species (Steiger et al., 2013). Then, we calculated the proportion of time points assigned as active in two time windows, between 10:00h to 16:00h (i.e., diurnal phase) and between 22:00h to 02:00h (i.e., nocturnal phase). Like in previous studies (e.g., Dominoni et al., 2014), we used fixed time windows to quantify activity levels to avoid biases introduced by seasonal changes in day length and individual times of onset and end of activity. The rationale to choose these windows was to have as much as possible of the relevant window (morning or evening) without risking including times when some birds might have already started or ended their activity (e.g., for the diurnal phase, we wanted not to include time in the morning when birds might not have started their activity yet, or time in the evening when birds might have ended their activity already).

### Environmental data

Interpolated daily minimum temperature and daily total precipitation were extracted from the MET Office HadUK dataset, which are available at a resolution of 1 x 1 km across the UK (Hollis et al., 2019). For each site included in this study, daily climatic variables were extracted from the central location of each site (see coordinates above).

### Statistical analysis

Bayesian models were fitted using the brms R package (v2.19.0; Bürkner, 2017). We used default priors. MCMC chain convergence was assessed visually and by computing R-hat values. We computed predictions and marginal effects using posterior distributions, calculating median and 95% Credible Intervals (‘95%CrI’ hereafter). We inferred statistical evidence for effects when 95%CrI did not include zero.

#### Onset of daily activity, end of daily activity and duration of diurnal activity

Variation in relative onset of activity, relative end of activity and duration of diurnal activity of all six species were analysed using three Bayesian linear mixed models (LMM), one for each response variable. In these three models, we included the same fixed effect structure. Models contained as fixed effects: habitat (2-level factor, ‘urban’ and ‘forest’), species (6-level factor), time of the year (2-level factor, ‘pre-breeding’ and ‘post-breeding’), year (2-level factor, ‘2020’ and ‘2021’), daily minimum temperature and daily precipitation. Sex is not included in the overarching model because males and females were identified in only three species (see below). To allow for differential urban effects across species and across times of the year, we included the 2-way interactions between habitat and species, and habitat and time of the year. We included these interactions as they directly test our predictions (see Introduction). Site of sampling and date of observation were included as random effect intercepts. Patterns of variation in relative time of onset and end of activity were similar between sites of sampling within habitat types (Figure S4; Figure S5). To allow for differences in between and within-individual variation across habitats, these models included habitat-specific bird ID random intercepts (i.e., modelling between-individual variation for each habitat) and habitat-specific residual variance terms (i.e., modelling within-individual variation for each habitat). These models were fitted running three MCMC chains of 50,000 iterations each including 20,000 warm-up iterations, which were discarded, and sampling every 10 iterations thereafter. To investigate whether urban conditions affect diel activity of both sexes equally, we re-ran the onset and end models described above for blackbirds, great tits and chaffinches, species that can be reliably sexed based on plumage traits, with the addition of sex and the interactions between sex and habitat, sex and time of the year, and sex and species as fixed effects (these models were fitted running three MCMC chains of 75,000 iterations each including 50,000 warm-up iterations, and sampling every 10 iterations thereafter).

#### Diurnal and nocturnal level of activity

We used a generalised linear mixed model (GLMM) with a binomial error structure to analyse the proportion of time that individuals were classified as active (i.e., activity level) during the diurnal phase (i.e., from 10:00h to 16:00h). We used that same type of GLMM to also analyse the proportion of time that individuals were classified as active during the nocturnal phase (i.e., from 22:00h to 02:00h). We first ran these two GLMMs separately, including the same fixed and random effect structure specified in the onset/end/duration models. As standard binomial GLMMs do not include residual variation, and thus within-individual variation cannot be calculated, we included a habitat-specific observation-level random effect that allowed us to model within-individual variation in activity levels (i.e., residual variation). We then combined the two response terms (i.e., diurnal and nocturnal proportion of activity) in one single bivariate binomial model. This model included the same fixed effect and random effect structure specified above for each of the single-response models, but it allowed us to model within and between-individual correlations of diurnal and nocturnal levels of activity; hence, effectively testing for the existence of the hypothesised trade-off between nocturnal and diurnal levels of activity. For these analyses, we included only observations (either diurnal or nocturnal) where individual data were available for at least 50% of the total diurnal or nocturnal window length (i.e., that an individual had been recorded at least 50% of the total time relevant for a given time window). These binomial GLMMs were fitted running three MCMC chains of 100,000 iterations each including 25,000 warm-up iterations and sampling every 50 iterations thereafter.

#### Within and between-individual variation, and repeatability, in activity traits

The models presented above provided us with estimations of between-individual variances for urban and forest populations. We used these to calculate habitat-specific repeatability for relative onset and end of daily activity, and for duration of diurnal activity. Repeatability was calculated by dividing between-individual variation by total phenotypic variation (i.e., proportion of the total variance accounted for by differences between individuals; (Nakagawa & Schielzeth, 2010). The use of full posterior distributions to make these calculations allowed us to directly compute 95%CrI for repeatability values.

## Results

### Relative onset of daily activity

We obtained data for onset of activity of 262 individuals, 149 individuals in the urban habitat and 113 individuals in the forest, from six species, totalling 3,855 observations. On average, activity started 0.46 hours before sunrise in urban habitats (standard deviation [SD] = 1.08 hours) and 0.13 hours before sunrise in forest habitats (SD = 0.78 hours) (Figure 1a). Urban effects on onset of activity varied among species (Figure 1a; Figure S6a; Table S2). Urban robins and blackbirds started their activity, on average, 30 and 33 minutes earlier, respectively, than their forest counterparts (Figure S6a). The onset of activity of urban great tits, blue tits, dunnock and chaffinch did not differ from their forest counterparts (Figure S6a). Urban effects on onset of activity did not vary across times of the year. In both seasons, pre- and post-breeding, urban effects were consistently negative (Figure S6b), with the formal comparison of these seasonal effects not yielding a significant difference (Figure S6b). Daily minimum temperatures and rainfall were not associated with time of onset of activity (Table S2). Urban effects on relative onset of activity did not differ between males and females in the three species for which data were available (Figure S7a).

**Figure 1.**
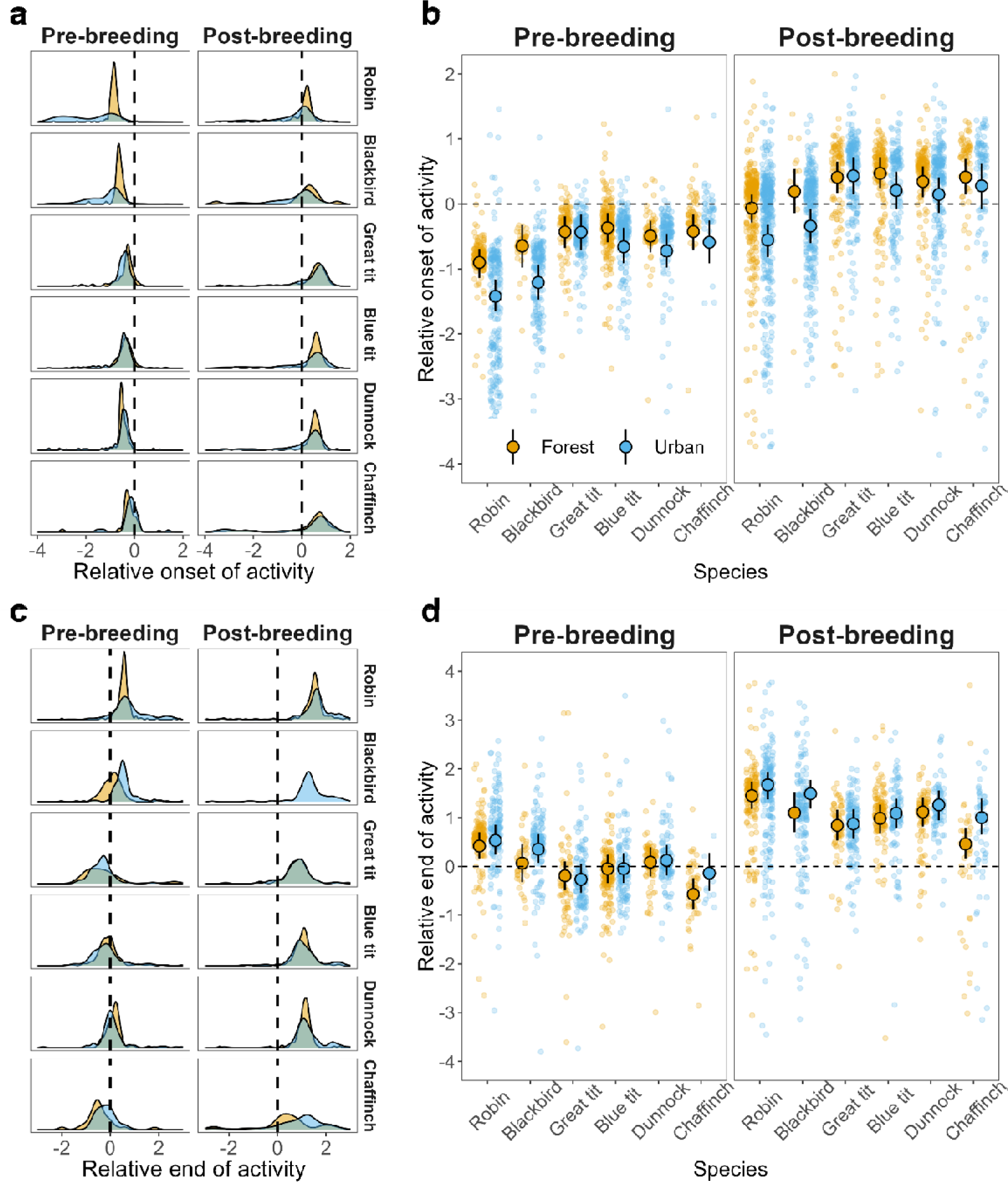
Onset and end of activity in urban and forest habitats post- and pre-breeding across passerine species. (**a**) Onset: Density plot of raw data for relative onset of activity in urban and forest habitats pre- and post-breeding across the six species. (**b**) Time of onset of activity relative to sunrise in urban and forest habitats pre- and post-breeding across the six species under study. (**c**) End: Density plot of raw data for relative end of activity in urban and forest habitats pre- and post-breeding across the six species. (**d**) Time of end of activity relative to sunset in urban and forest habitats in pre- and post-breeding across the six species. In (b) and (d), dashed lines represent sunrise time and sunset time, respectively, and the raw data are shown as small translucid points with filled points and intervals illustrating model predictions and 95% Credible Interval (95%CrI). Blue shade indicates urban populations, while yellow shade indicates forest populations.

Variation in time of onset of activity was higher in urban birds than in forest birds (Table S2; Figure 2a), both between individuals (posterior estimate of the difference in between-individual standard deviation in urban and forest habitats [95%CrI] = 0.102 [+0.020, +0.188]), and within individuals (posterior estimate of the difference in within-individual [i.e., residual] standard deviation in urban and forest habitats [95%CrI] = 0.259 [+0.225, +0.292]). Repeatability of relative onset of activity was similar across habitats (estimate [95%CrI] for forest birds = 0.169 [0.104, 0.234]; estimate [95%CrI] for urban birds = 0.159 [0.112, 0.213]); these repeatability estimates did not differ from each other (posterior difference estimate [95%CrI] = −0.008 [−0.088, +0.063]).

**Figure 2.**
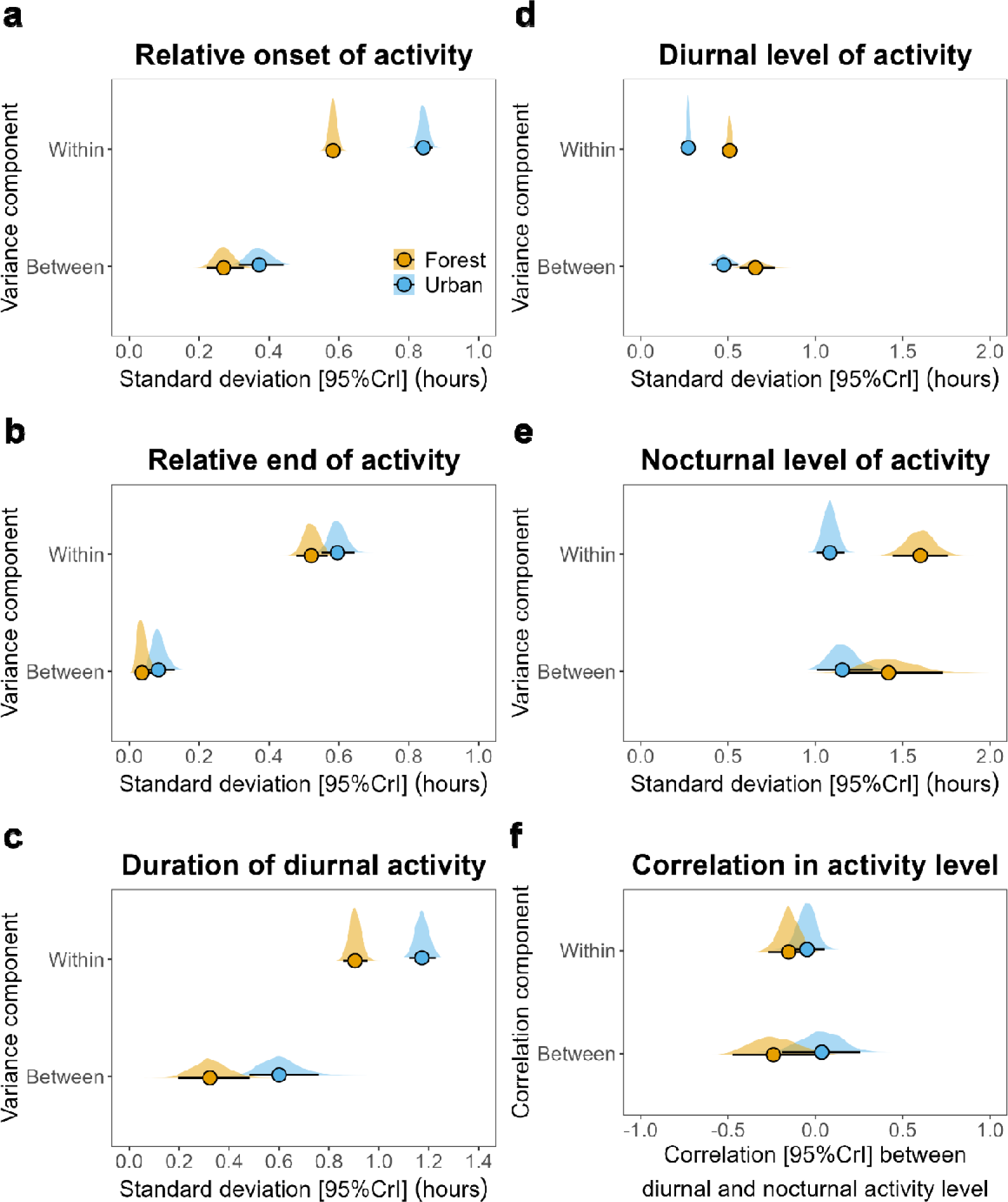
Within and between-individual variance components in urban and forest habitats. Variance components (**a**) for time of relative onset of activity, (**b**) time of relative end of activity, (**c**) duration of diurnal activity, (**d**) levels of diurnal activity and (**e**) level of nocturnal activity. (**f**) Correlation between nocturnal and diurnal level of activity between and within individuals. Posterior median and 95% Credible Intervals [95% CrI] for variance and correlation components (variances presented as model estimated standard deviations) given along with the full posterior distribution for urban (blue) and forest (orange) habitats.

### Relative end of daily activity

We obtained data for end of activity of 213 individuals, 117 in the urban habitat and 96 in the forest, from six species totalling 2,601 observations. On average, activity ended 0.74 hours after sunset in urban habitats (SD = 1.04 hours) and 0.56 hours before sunset in forest habitats (SD = 0.95 hours) (Figure 1c). Urban effects on end of activity varied among species (Figure 1c; Figure S6c; Table S3), with chaffinch and blackbirds showing larger shifts in end of activity (Figure S6c); only chaffinch showed significant differences in end of activity between urban and forest habitats, with urban chaffinch ending their activity later than forest chaffinch (Figure S6c). Urban effects on end of activity did not vary across times of the year (Figure S6d). Daily minimum temperatures negatively predicted end of activity: the colder a given day, the later the end of activity (Table S3). Daily rainfall was not associated with time of end of activity (Table S3). Urban effects on relative end of activity did not differ between males and females (Figure S7b).

Variation in time of end of activity was higher in urban birds than in forest birds (Table S3; Figure 2b), in particular within individuals (posterior estimate of the difference in between-individual standard deviation in urban and forest habitats [95%CrI] = 0.099 [−0.004, +0.201]; posterior estimate of the difference in within-individual [i.e., residual] standard deviation in urban and forest habitats [95%CrI] = 0.050 [+0.008, +0.094]). Despite higher within-individual variation, repeatability of relative end of activity was higher in urban birds (estimate [95%CrI] = 0.115 [0.069, 0.172]) than in forest birds (estimate [95%CrI] = 0.060 [0.023, 0.120]); these repeatability estimates did not statistically differ from each other (posterior difference estimate [95%CrI] = 0.0.053 [−0.016, +0.122]).

### Duration of diurnal activity

Combining daily estimates of onset and end of activity, we estimated duration of daily activity (i.e., time elapsed between the onset and the end of activity of a given day) for 195 individuals of five species with a total of 1,968 daily observations. The estimated mean duration of diurnal activity was 12.2 hours (SD = 1.7 hours) for urban birds and 11.7 hours (SD = 1.6 hours) for forest birds. Urban effects on the duration of diurnal activity varied among species (Table S4; Figure S8a), with chaffinch and robin showing the largest habitat-associated differences (Figure S8b); however, no species showed significant differences in duration of activity between urban and forest habitats (Figure S8b). Urban effects on duration of activity did not vary across times of the year (Figure S8c). Overall, urban – forest differences in duration of activity showed large credible intervals which included zero in both the pre-breeding and post-breeding phases, and did not differ from each other (Figure S8c). Daily rainfall did not predict duration of activity (Table S4). Duration of activity was shorter in 2021 compared to 2020 (Table S4) and daily minimum temperatures were positively associated with duration of diurnal activity (Table S4).

Variation in duration of activity was higher in urban birds than in forest birds (Table S4; Figure 2c), both between individuals (posterior estimate of the difference in between-individual standard deviation in urban and forest habitats [95%CrI] = 0.278 [+0.076, +0.472]) and within individuals (posterior estimate of the difference in within-individual [i.e., residual] standard deviation in urban and forest habitats [95%CrI] = 0.269 [+0.196, +0.342]). Repeatability of duration of activity was higher in urban birds (estimate [95%CrI] = 0.149 [0.080, 0.226]) than in forest birds (estimate [95%CrI] = 0.063 [0.022, 0.145]); these repeatability estimates differed from each other (posterior difference estimate [95%CrI] = 0.081 [+0.001, +0.167]).

### Diurnal and nocturnal levels of activity

We obtained data to calculate diurnal activity levels for 274 individuals, 147 in the urban habitat and 127 in the forest, totalling 3,785 observations. Overall, urban birds were active 48.5% of their diurnal phase (SD = 11.7%); forest birds were active 49.7% of their day (SD = 15.4%). Diurnal activity levels varied between species (Table S5; Figure S9a-b), with great tit, blue tit and robin showing the highest levels of activity (Figure S9a). Urban and forest populations had similar levels of diurnal activity for all species (Figure S9b). There were no differences in levels of diurnal activity between urban and forest birds in either the pre- or post-breeding time of the year (Figure S9c). Year of study, daily minimum temperatures and rainfall were not associated with diurnal levels of activity (Table S5). Variation in diurnal levels of activity was lower in urban birds than in forest birds (Table S5; Figure 2d), both between individuals (posterior estimate of the difference in between-individual standard deviation in urban and forest habitats [95%CrI] = −0.182 [−0.312, −0.056]) and within individuals (posterior estimate of the difference in within-individual [i.e., residual] standard deviation in urban and forest habitats [95%CrI] = −0.237 [−0.265, −209]).

We obtained data to calculate nocturnal activity levels for 300 individuals, 176 in the urban habitat and 124 in the forest, amounting to 5,106 observations. Overall, urban birds were active 4.1% of the nocturnal phase (SD = 9.6%); forest birds were active 1.6% of the nocturnal period (SD = 4.9 %). Across all species, urban populations displayed higher levels of nocturnal activity than forest populations (Figure S10b), and this effect was particularly strong for dunnock (48.8% increase in the urban habitat), blackbird (42.4% increase in the urban habitat) and robin (36.0% increase in the urban habitat; Figure 10b). Urban populations displayed higher levels of nocturnal activity in both pre- and post-breeding phases (Figure S10c), with a particularly strong effect in the post-breeding phase (Figure S10c). Year of study, daily minimum temperatures and rainfall were not associated with nocturnal levels of activity (Table S6). Within-individual variation in nocturnal levels of activity was lower in urban birds than in forest birds (Figure 2e; Table S6). Between-individual variation in nocturnal levels of activity did not differ across habitats (Figure 2e; Table S6).

After assessing urban effects on the variation and mean of diurnal and nocturnal activity levels separately, we combined both models in a single bivariate model (i.e., having two response variables) to quantify whether individuals compensate during the day for increased nocturnal activity. 5,871 observations with information for either or both nocturnal and diurnal levels of activity were included in this model. We did not find statistical evidence of negative (as predicted) or positive correlations between diurnal and nocturnal activity levels in the urban or forest habitat at the between-individual level (i.e., correlation values different from zero; Table S7, Figure 2f); however, diurnal and nocturnal levels of activity were negatively correlated at the within-individual level in the forest, but not in the urban habitat (Table S7, Figure 2f). The magnitude of between and within-individual correlations in diurnal and nocturnal activity levels did not significantly differ across habitats (posterior estimate of the difference in diurnal and nocturnal activity level correlations across habitats: between individuals [95%CrI] = 0.273 [−0.063, 0.609]; within individuals [95%CrI] = 0.104 [−0.044, 0.251]).

## Discussion

To test how urban conditions affect activity patterns across a range of passerine species, we carried out a concurrent, comparative and in-depth automated telemetry study, including individuals from two urban and two forest populations over two years. The analysis of this dataset shows that urban birds have modified patterns of activity with, overall, earlier onset of activity, higher levels of nocturnal activity and, more between- and within-individual variation in onset and end of activity than forest birds. Additionally, our findings reveal that the (correlative) effects of urban conditions on diel patterns of activity vary across species in a community of passerine birds and that they are consistent across sexes.

As expected, early-rising species showed the strongest shifts in diel patterns of activity in urban conditions. Urban blackbirds (both males and females) and robins showed earlier onset of activity than their forest counterparts. Such behavioural differences have been described for male blackbirds (e.g., Dominoni et al., 2013) and for incubating female great tits (Tomotani et al., 2023; Womack et al., 2023), but, it was not clear if these patterns would exist in other species (e.g., robins). We did not find a statistically significant effect of urban conditions on time of onset of activity of blue tits and dunnock. However, it is worth nothing that our data indicates a clear tendency for these species to start their activity earlier in the urban habitat than in the forest. Urban great tits and chaffinches started their daily activity at similar times as their forest counterparts. These results for behavioural patterns of activity contrast with previous attempts to quantify the effects of urban factors on timing of singing (Kempenaers et al., 2010; Silva et al., 2014). In that case, blackbirds, robins, but also great tits and blue tits exposed to artificial light at night were found to start singing earlier than counterparts in dark habitats (Kempenaers et al., 2010; Silva et al., 2014). Both sets of results are not mutually exclusive though, as great tits and blue tits might start singing earlier in urban habitats while still in their night roosts (i.e., before the onset of activity). Interestingly, the effects of ALAN on the timing of singing (Silva et al., 2014) and the urban effects on activity patterns reported here ranked similarly across species, with the strongest effect on blackbird and robin, then on great tit and blue tit, and then on chaffinch (for which no differences were found for time of singing onset or patterns of activity in da Silva et al., 2014). While the mechanism explaining these species-specific responses to urban conditions is not yet clear, it has been suggested that the natural sensitivity of species to low light conditions plays a role (McNeil et al., 2005; Thomas et al., 2002), with highly light-sensitive species showing stronger shifts in diel behaviour (e.g., blackbird and robin) than less light-sensitive species (e.g., chaffinch and dunnock). In addition or alternatively, the different responses may be linked with dietary specialisations (Morelli et al., 2023). As evidenced in previous studies, urban effects on the end of activity were much weaker compared to the effects on the onset of activity. While our data does not provide a mechanism for this contrast between urban effects on onset and end of activity, higher levels of selection are expected in traits that are expressed in the morning (Hau et al., 2017), perhaps pre-disposing these traits to stronger urban effects.

Our study also reveals that urban effects on diel activity patterns vary throughout the year. First, the advancement of activity in urban birds was larger in the pre-breeding phase than in the post-breeding phase. This was in line with our prediction that if early rising has reproductive advantages, then we should expect to see stronger differences between urban and forest birds during the breeding season, or right before it when birds are preparing to breed (i.e., during the phase of gonadal growth that boosts the production of steroid hormones and during mating; Gwinner, 1975). Second, while urban birds showed, on average, higher levels of nocturnal activity than their forest counterparts, the increase in urban nocturnal activity was particularly high in the post-breeding phase compared to the pre-breeding phase. Interestingly, while nocturnal activity levels were increased in the urban environment, we did not find evidence of carry-over effects into the day. We predicted that high levels of nocturnal activity would be correlated with lower levels of diurnal activity, particularly in the urban environment where birds are more active at night. However, we found an indication of this negative correlation in forest but not in urban habitats: there was a negative correlation between levels of diurnal and nocturnal activity at the individual level in the forest but not in the urban habitat. This might mean that forest birds have the ability, or else the need, to compensate for higher activity at night by resting more during the day. Conversely, urban birds could be prevented from daytime resting by human disturbance and traffic noise (Connelly et al., 2020), or else they might be better able to sustain higher levels of activity throughout the whole 24 hours. A similar, but not statistically significant, trend of (negative) correlation between diurnal and nocturnal levels of activity was found at the between individual level in the forest, suggesting temporal niche partition between individuals (i.e., with individuals that consistently show high levels of activity during the day having low levels of activity at night and *vice versa*).

Our analyses further reveal strong urban effects on the level of between and within-individual variation in time of onset of activity, time of end of activity and duration of diurnal activity: for these traits, urban individuals were behaviourally less synchronous with each other (i.e., higher between-individual variation in the urban habitat than in the forest) and also behaviourally less consistent (i.e., higher within-individual variation in the urban habitat than in the forest). It has recently been theorised that urban conditions could increase phenotypic variation (Thompson et al., 2022). Such patterns have been documented for bird phenology (CapillalJLasheras et al., 2022) and are also starting to emerge for morphological and behavioural traits (Dominoni et al., 2014; Thompson et al., 2022). Previous attempts at quantifying differences in phenotypic variation between urban and non-urban populations could not, however, disentangle the sources of variation that generated the observed increase of variation in urban populations (CapillalJLasheras et al., 2022). Given our ability to quantify both between and within-individual variation in traits associated with patterns of activity, our findings might suggest that both higher genotypic variation (i.e., higher among individual variation) and higher level of plasticity (i.e., higher within-individual variation) exist in urban populations, at least in connection to the time-keeping traits investigated here. Higher habitat heterogeneity (CapillalJLasheras et al., 2022; Morelli et al., 2023), carry-over effects due to habituation to microenvironmental conditions (e.g., ALAN) or higher variation in direct human disturbance in urban habitats (Gaynor et al., 2018; Geraghty & O’Mahony, 2016) could cause higher within-individual variation through plasticity in the urban habitat. This does not necessarily mean that urban populations are more plastic than non-urban populations *per se*, but simply that they are exposed to a more variable set of environmental conditions (Baythavong, 2011; Gomez-Mestre & Jovani, 2013). Interestingly, despite higher (between and within-individual) phenotypic variation in timing traits in urban populations, variation in diurnal and nocturnal activity levels tended to be lower in urban habitats. We currently have no explanation for the observation that activity levels, both during the day and night, of forest birds are more variable than activity levels of urban birds. This result, however, highlights that the hypothesised increase in phenotypic variation in urban habitats does not apply to all phenotypic traits. These results also emphasise that a mechanistic understanding of how urban conditions affect phenotypic variation is essential to predict trait-specific responses to urbanisation, fitness consequences and their eco-evolutionary effects (Senzaki et al., 2020).

## Conclusions

Using automated telemetry in multiple urban and forest populations of six passerine birds to record their activity patterns, we have shown that the association between urbanisation and changes in patterns of activity varies across species, possibly due to species differences in sensitivity to low-lighting conditions. Importantly, these shifts in activity patterns of urban populations are dependent on time of the year, with pre-breeding urban effects on time of onset of activity being stronger than post-breeding urban effects. Conversely, increases in nocturnal activity associated with the urban habitat were much stronger in the post-breeding phase compared to the pre-breeding phase. Urban populations display higher phenotypic variation than forest populations, as reported here for activity timing but not for activity levels contributes to our understanding of the mechanisms of adjustment to urban conditions. Our results also have methodological implications. For example, statistical models used in ecological research often assume constant variance across groups of observations. This assumption might often be violated when comparing urban *versus* non-urban animal populations. Current modelling techniques, however, can include predictors on variance components, relaxing this assumption and potentially fully revealing urban effects on phenotypic means and variation. Because urban effects on patterns of bird activity vary between species and throughout the year, nuanced approaches are required to fully understand the extent to which urban-induced changes in activity rhythms might be adaptive and/or have consequences for animal health.

## Acknowledgements

We thank Alistair Laming, Crinan Jarrett, Hannah Greetham, Hannah Humphreys, Jennifer Page, Mark Pitt, Paul Noyes, Phoebe Owen for help during field work. We thank the staff of SCENE, particularly Matt Newton and Hannele Honkanen, for hosting us and supporting our work. We thank Glasgow City Council and the Loch Lomond and the Trossachs National Park for giving us access to the urban and forest field sites, respectively. This research was funded by a NERC Highlight Topics grant to DMD (grant number: NE/S005773/1). PC-L is also funded by the European Union’s Horizon Europe research and innovation programme under the Marie Skłodowska-Curie grant agreement No 101150591.

## Author contributions

DMD, BH, PC-L and CJB designed the study. DMD, PC-L, CJB and PJB collected the data. PC-L and CC analysed the data. PC-L and DMD wrote the article, with input from all co-authors.

## Statement of inclusion

All authors were based in the country where the study was carried out.

## Data availability statement

All R scripts and datasets needed to reproduce the analyses presented in this paper are available at: https://github.com/PabloCapilla/variation_activity. Should the manuscript be accepted, a DOI to this data repository will be provided.

## References

1. Aulsebrook, A. E., Lesku, J. A., Mulder, R. A., Goymann, W., Vyssotski, A. L., & Jones, T. M. (2020). Streetlights Disrupt Night-Time Sleep in Urban Black Swans. Frontiers in Ecology and Evolution, 8, 131. 10.3389/fevo.2020.00131

2. Baythavong, B. S. (2011). Linking the spatial scale of environmental variation and the evolution of phenotypic plasticity: Selection favors adaptive plasticity in fine-grained environments. American Naturalist, 178, 75–87. 10.1086/660281

3. Branston, C. J., Capilla-Lasheras, P., Pollock, C. J., Griffiths, K., White, S., & Dominoni, D. M. (2021). Urbanisation weakens selection on the timing of breeding and clutch size in blue tits but not in great tits. Behavioral Ecology and Sociobiology, 75(11), 155. 10.1007/s00265-021-03096-z

4. Bürkner, P.-C. (2017). brms: An R Package for Bayesian Multilevel Models Using Stan. Journal of Statistical Software, 10(1), 1–28. 10.18637/jss.v080.i01

5. Capilla-Lasheras, P., Dominoni, D. M., Babayan, S. A., O’Shaughnessy, P. J., Mladenova, M., Woodford, L., Pollock, C. J., Barr, T., Baldini, F., & Helm, B. (2017). Elevated Immune Gene Expression Is Associated with Poor Reproductive Success of Urban Blue Tits. Frontiers in Ecology and Evolution, 5, 64. 10.3389/fevo.2017.00064

6. Capilla-Lasheras, P., Thompson, M. J., Sánchez-Tójar, A., Haddou, Y., Branston, C. J., Réale, D., Charmantier, A., & Dominoni, D. M. (2022). A global meta-analysis reveals higher variation in breeding phenology in urban birds than in their non-urban neighbours. Ecology Letters, 25(11), 2552–2570. 10.1111/ele.14099

7. Connelly, F., Johnsson, R. D., Aulsebrook, A. E., Mulder, R. A., Hall, M. L., Vyssotski, A. L., & Lesku, J. A. (2020). Urban noise restricts, fragments, and lightens sleep in Australian magpies. Environmental Pollution, 267, 115484. 10.1016/j.envpol.2020.115484

8. Daan, S., & Aschoff, J. (1975). Circadian rhythms of locomotor activity in captive birds and mammals: Their variations with season and latitude. Oecologia, 18, 269–316. 10.1007/BF00345851

9. De Framond, L., & Brumm, H. (2022). Long-term effects of noise pollution on the avian dawn chorus: A natural experiment facilitated by the closure of an international airport. Proceedings of the Royal Society B: Biological Sciences, 289(1982), 20220906. 10.1098/rspb.2022.0906

10. Dominoni, D. M., Carmona-Wagner, E. O., Hofmann, M., Kranstauber, B., & Partecke, J. (2014). Individual-based measurements of light intensity provide new insights into the effects of artificial light at night on daily rhythms of urban-dwelling songbirds. Journal of Animal Ecology, 83(3), 681–692. 10.1111/1365-2656.12150

11. Dominoni, D. M., De Jong, M., Van Oers, K., O’Shaughnessy, P., Blackburn, G. J., Atema, E., Mateman, A. C., D’Amelio, P. B., Trost, L., Bellingham, M., Clark, J., Visser, M. E., & Helm, B. (2022). Integrated molecular and behavioural data reveal deep circadian disruption in response to artificial light at night in male Great tits (Parus major). Scientific Reports, 12(1), 1553. 10.1038/s41598-022-05059-4

12. Dominoni, D. M., Halfwerk, W., Baird, E., Buxton, R. T., Fernández-Juricic, E., Fristrup, K. M., McKenna, M. F., Mennitt, D. J., Perkin, E. K., Seymoure, B. M., Stoner, D. C., Tennessen, J. B., Toth, C. A., Tyrrell, L. P., Wilson, A., Francis, C. D., Carter, N. H., & Barber, J. R. (2020). Why conservation biology can benefit from sensory ecology. Nature Ecology and Evolution, 4, 502–511. 10.1038/s41559-020-1135-4

13. Dominoni, D. M., Helm, B., Lehmann, M., Dowse, H. B., & Partecke, J. (2013). Clocks for the city: Circadian differences between forest and city songbirds. Proceedings of the Royal Society B, 280(1763), 20130593. 10.1098/rspb.2013.0593

14. Foster, R. G., & Kreitzman, L. (2005). Rhythms of Life. Yale University Press.

15. Gaynor, K. M., Hojnowski, C. E., Carter, N. H., & Brashares, J. S. (2018). The influence of human disturbance on wildlife nocturnality. Science, 360, 1232–1235. 10.1126/science.aar7121

16. Geraghty, D., & O’Mahony, M. (2016). Investigating the temporal variability of noise in an urban environment. International Journal of Sustainable Built Environment, 5(1), 34– 45.

17. Gilbert, N. A., McGinn, K. A., Nunes, L. A., Shipley, A. A., Bernath-Plaisted, J., Clare, J. D. J., Murphy, P. W., Keyser, S. R., Thompson, K. L., Maresh Nelson, S. B., Cohen, J. M., Widick, I. V., Bartel, S. L., Orrock, J. L., & Zuckerberg, B. (2023). Daily activity timing in the Anthropocene. Trends in Ecology & Evolution, 38(4), 324–336. 10.1016/j.tree.2022.10.008

18. Gomez-Mestre, I., & Jovani, R. (2013). A heuristic model on the role of plasticity in adaptive evolution: Plasticity increases adaptation, population viability and genetic variation. Proceedings of the Royal Society B, 280, 20131869. 10.1098/rspb.2013.1869

19. Gwinner, E. (1975). Die circannuale Periodik der Fortpflanzungsaktivität beim Star (Sturnus vulgaris) unter dem Einfluß gleich- und andersgeschlechtiger Artgenossen. Zeitschrift Für Tierpsychologie, 38(1), 34–43. 10.1111/j.1439-0310.1975.tb01990.x

20. Hasselt, S. J., Hut, R. A., Allocca, G., Vyssotski, A. L., Piersma, T., Rattenborg, N. C., & Meerlo, P. (2021). Cloud cover amplifies the sleep-suppressing effect of artificial light at night in geese. Environmental Pollution, 273, 116444. 10.1016/j.envpol.2021.116444

21. Hau, M., Dominoni, D. M., Casagrande, S., Buck, C. L., Wagner, G., Hazlerigg, D., Greives, T., & Hut, R. A. (2017). Timing as a sexually selected trait: The right mate at the right moment. Philosophical Transactions of the Royal Society, 372, 20160249. 10.1098/rstb.2016.0249

22. Helm, B., Visser, M. E., Schwartz, W., Kronfeld-Schor, N., Gerkema, M., Piersma, T., & Bloch, G. (2017). Two sides of a coin: Ecological and chronobiological perspectives of timing in the wild. Philosophical Transactions of the Royal Society B, 372, 20160246. 10.1098/rstb.2016.0246

23. Hollis, D., McCarthy, M., Kendon, M., Legg, T., & Simpson, I. (2019). HadUK-Grid—A new UK dataset of gridded climate observations. Geoscience Data Journal, 6(2), 151–159. 10.1002/gdj3.78

24. Jarrett, C., Powell, L. L., McDevitt, H., Helm, B., & Welch, A. J. (2020). Bitter fruits of hard labour: Diet metabarcoding and telemetry reveal that urban songbirds travel further for lower-quality food. Oecologia, 193(2), 377–388. 10.1007/s00442-020-04678-w

25. Kacelnik, A., & Krebs, J. R. (1983). The Dawn Chorus in the Great Tit (Parus Major): Proximate and Ultimate Causes…Behaviour, 83, 287–308.

26. Kempenaers, B., Borgström, P., Loës, P., Schlicht, E., & Valcu, M. (2010). Artificial night lighting affects dawn song, extra-pair siring success, and lay date in songbirds. Current Biology, 20, 1735–1739. 10.1016/j.cub.2010.08.028

27. Kronfeld-Schor, N., & Dayan, T. (2003). Partitioning of Time as an Ecological Resource. Annual Review of Ecology, Evolution, and Systematics, 34, 153–181. 10.1146/annurev.ecolsys.34.011802.132435

28. Martinez-Bakker, M., & Helm, B. (2015). The influence of biological rhythms on host– parasite interactions. Trends in Ecology & Evolution, 30(6), 314–326. 10.1016/j.tree.2015.03.012

29. Maury, C., Serota, M. W., & Williams, T. D. (2020). Plasticity in diurnal activity and temporal phenotype during parental care in European starlings, Sturnus vulgaris. Animal Behaviour, 159, 37–45. 10.1016/j.anbehav.2019.11.004

30. McGlade, C. L. O., Capilla-Lasheras, P., Womack, R. J., Helm, B., & Dominoni, D. M. (2023). Experimental light at night explains differences in activity onset between urban and forest great tits. Biology Letters, 19(9), 20230194. 10.1098/rsbl.2023.0194

31. McNeil, R., McSween, A., & Lachapelle, P. (2005). Comparison of the retinal structure and function in four bird species as a function of the time they start singing in the morning. Brain, Behavior and Evolution, 65(3), 202–214. 10.1159/000083881

32. Morelli, F., Tryjanowski, P., Ibáñez-Álamo, J. D., Díaz, M., Suhonen, J., Pape Møller, A., Prosek, J., Moravec, D., Bussière, R., Mägi, M., Kominos, T., Galanaki, A., Bukas, N., Markó, G., Pruscini, F., Reif, J., & Benedetti, Y. (2023). Effects of light and noise pollution on avian communities of European cities are correlated with the species’ diet. Scientific Reports, 13(1), 4361. 10.1038/s41598-023-31337-w

33. Nakagawa, S., & Schielzeth, H. (2010). Repeatability for Gaussian and non-Gaussian data: A practical guide for biologists. Biological Reviews, 85(4), 935–956. 10.1111/j.1469-185X.2010.00141.x

34. Ouyang, J. Q., Jong, M., Grunsven, R. H. A., Matson, K. D., Haussmann, M. F., Meerlo, P., Visser, M. E., & Spoelstra, K. (2017). Restless roosts: Light pollution affects behavior, sleep, and physiology in a free-living songbird. Global Change Biology, 23, 4987– 4994. 10.1111/gcb.13756

35. Poesel, A., Kunc, H. P., Foerster, K., Johnsen, A., & Kempenaers, B. (2006). Early birds are sexy: Male age, dawn song and extrapair paternity in blue tits, Cyanistes (formerly Parus) caeruleus. Animal Behaviour, 72(3), 531–538. 10.1016/j.anbehav.2005.10.022

36. Pollock, C. J., Capilla-Lasheras, P., McGill, R. A. R., Helm, B., & Dominoni, D. M. (2017). Integrated behavioural and stable isotope data reveal altered diet linked to low breeding success in urban-dwelling blue tits (Cyanistes caeruleus. Scientific Reports, 7, 5014. 10.1038/s41598-017-04575-y

37. Raap, T., Pinxten, R., & Eens, M. (2015). Light pollution disrupts sleep in free-living animals. Scientific Reports, 5. 10.1038/srep13557

38. Sanders, D., Frago, E., Kehoe, R., Patterson, C., & Gaston, K. J. (2020). A meta-analysis of biological impacts of artificial light at night. Nature Ecology & Evolution, 5(1), 74–81. 10.1038/s41559-020-01322-x

39. Senzaki, M., Barber, J. R., Phillips, J. N., Carter, N. H., Cooper, C. B., Ditmer, M. A., Fristrup, K. M., McClure, C. J. W., Mennitt, D. J., Tyrrell, L. P., Vukomanovic, J., Wilson, A. A., & Francis, C. D. (2020). Sensory pollutants alter bird phenology and fitness across a continent. Nature, 587(7835), 605–609. 10.1038/s41586-020-2903-7

40. Serota, M. W., & Williams, T. D. (2019). Adjustment of total activity as a response to handicapping European starlings during parental care. Animal Behaviour, 148, 19–27. 10.1016/j.anbehav.2018.11.009

41. Silva, A., Samplonius, J. M., Schlicht, E., Valcu, M., & Kempenaers, B. (2014). Artificial night lighting rather than traffic noise affects the daily timing of dawn and dusk singing in common European songbirds. Behavioral Ecology, 25, 1037–1047. 10.1093/beheco/aru103

42. Silva, A., Valcu, M., & Kempenaers, B. (2015). Light pollution alters the phenology of dawn and dusk singing in common European songbirds. Philosophical Transactions of the Royal Society of London. Series B, Biological Sciences, 370(1667), 20140126-. 10.1098/rstb.2014.0126

43. Steiger, S. S., Valcu, M., Spoelstra, K., Helm, B., Wikelski, M., & Kempenaers, B. (2013). When the sun never sets: Diverse activity rhythms under continuous daylight in free-living arctic-breeding birds. Proceedings of the Royal Society B, 280(1764), 20131016. 10.1098/rspb.2013.1016

44. Sun, J., Raap, T., Pinxten, R., & Eens, M. (2017). Artificial light at night affects sleep behaviour differently in two closely related songbird species. Environmental Pollution, 231, 882–889. 10.1016/j.envpol.2017.08.098

45. Thomas, R. J., Székely, T., Cuthill, I. C., Harper, D. G. C., Newson, S. E., Frayling, T. D., & Wallis, P. D. (2002). Eye size in birds and the timing of song at dawn. Proceedings of the Royal Society B, 269, 831–837. 10.1098/rspb.2001.1941

46. Thompson, M. J., Capilla-Lasheras, P., Dominoni, D. M., Réale, D., & Charmantier, A. (2022). Phenotypic variation in urban environments: Mechanisms and implications. Trends in Ecology & Evolution, 37(2), 171–182. 10.1016/j.tree.2021.09.009

47. Tomotani, B. M., Timpen, F., & Spoelstra, K. (2023). Ingrained city rhythms: Flexible activity timing but more persistent circadian pace in urban birds. Proceedings of the Royal Society B, 290, 20222605. 10.1098/rspb.2022.2605

48. Womack, R. J., Capilla-Lasheras, P., McGlade, C. L. O., Dominoni, D. M., & Helm, B. (2023). Reproductive fitness is associated with female chronotype in a songbird. Animal Behaviour, 205, 65–78. 10.1016/j.anbehav.2023.08.018

