## Supplementary materials for "Urban effects on timing and variability of diel activity differ across passerine species and seasons"

**Supplementary methods**

*1 - Bayesian broken stick approach to estimate onset and end of activity*

We validated the onset and end of activity estimates based on a frequentist change point model (Dominoni et al., 2013) using an alternative, independent, method for onset and end of activity estimation, a Bayesian Broken-Stick model fitted using the R package ‘mcp’ (v0.3.2; Lindeløv, 2020). This model fitted two regression lines connected via a breaking point (i.e., estimated time of onset or end of activity). The two regression sub-models were specified as:

$\left\{ \begin{aligned} y^{i}={Intercept}_{1}+\varepsilon_{1}^{2} if i<c \\ y^{i}={Intercept}_{2}+\varepsilon_{2}^{2} if i>c \end{aligned} \right.$(Eq. 1)

Where each $i$th observation was drawn from one of two normal distributions (subscripts ‘1’ and ‘2’ in Eq. 1) that were allowed to vary in their intercepts (‘$Intercept$’ in Eq. 1) and their residual variation (‘$\varepsilon$’ in Eq. 1). The model also calculated the value for ‘$c$’ (see in Eq. 1) which marks the breaking point between distribution ‘1’ and ‘2’. This model was fitted using three Markov Chain Monte Carlo (‘MCMC’) simulations of 4500 iterations discarding the first 1500 iterations. We assessed MCMC convergence for each model parameter calculating R-hat values. We considered that valid estimations of time of onset or end of activity had been reached when MCMCs had converged and showed an R-hat value lower than 1.01.

**Supplementary results**

*1 – Comparison of model results when time of onset and end of activity were estimated via frequentist versus Bayesian broken stick models*

Regardless of the method used to estimate onset and end of activity, our analyses led to very similar conclusions. The complementary Bayesian method to estimate time of onset and end of activity did produce similar results to the frequentist method (i.e., results shown in the main text). A full comparison of the results is provided in Table S2, Table S3 and Figure S11. In short, for the analysis of relative time of onset of activity, this complementary analysis also provided evidence for earlier relative onset of activity in urban robins and blackbirds compared to their forest counterparts (urban *versus* forest difference [95% Credible Intervals, CrI], for robins = -0.418 [-0.856, 0.034]; for blackbirds = -0.603 [-1.097, -0.073]) and similar pre- and post-breeding urban effects for all species (urban *versus* forest difference [95%CrI], for pre-breeding season = -0.210 [-0.532, 0.292]; for post-breeding season = -0.114 [-0.461, 0.304]; Table S2; Figure S11a). For the analysis of relative time of end of activity, the analysis of the complementary Bayesian estimation also led to similar conclusions to those presented in the main text, providing no evidence of urban effects on relative time of end of activity for any studied species and no evidence of general urban effects during the pre- and post-breeding seasons (Table S3; Figure S11b). We did not repeat the analysis of duration of diurnal activity after applying the complementary Bayesian method because duration of diurnal activity is directly computed from time of onset and end of activity (for which we show both approaches lead to similar conclusions).

**Supplementary Tables**

**Table S1.** Sample size used for the analysis of each response variable included in the main text per sex, species, year and habitat of sampling. ‘obs’ = number of observations, ‘ind’ = number of individuals. Values in brackets provide the number of individuals sampled in both times of the year (i.e., number of individuals sampled both in the pre- and post-breeding periods) and both years (i.e., number of individuals sampled both in 2020 and 2021). Overall, we equipped 6 blackbirds, 31 blue tits, 29 chaffinches, 22 dunnock, 35 great tits and 41 robins in forest habitats; and, 37 blackbirds, 33 blue tits, 17 chaffinches, 31 dunnock, 37 great tits and 47 robins in urban habitats.

|  |  | Time of onset of activity | | Time of end of activity | | Duration of diurnal activity | | Level of diurnal activity | | Level of nocturnal activity | |
| --- | --- | --- | --- | --- | --- | --- | --- | --- | --- | --- | --- |
|  |  | obs | ind | obs | ind | obs | ind | obs | ind | obs | ind |
| Habitat | Forest | 1592 | 113 | 1158 | 96 | 962 | 83 | 1767 | 127 | 1883 | 124 |
|  | Urban | 2263 | 149 | 1443 | 117 | 1006 | 86 | 2018 | 147 | 3223 | 176 |
| Sex | Female | 644 | 46 | 416 | 37 | 253 | 26 | 583 | 47 | 904 | 52 |
|  | Male | 731 | 56 | 475 | 43 | 222 | 19 | 712 | 65 | 1010 | 68 |
|  | Unknown | 2480 | 160 | 1710 | 133 | 1493 | 124 | 2490 | 162 | 3192 | 180 |
| Species | Blackbird | 540 | 34 | 332 | 29 | 0 | 0 | 431 | 33 | 793 | 41 |
|  | Blue tit | 730 | 45 | 532 | 40 | 474 | 38 | 839 | 54 | 796 | 49 |
|  | Chaffinch | 283 | 29 | 174 | 20 | 140 | 17 | 240 | 27 | 474 | 35 |
|  | Dunnock | 620 | 43 | 419 | 35 | 359 | 32 | 502 | 37 | 871 | 50 |
|  | Great tit | 583 | 41 | 410 | 33 | 356 | 30 | 653 | 55 | 681 | 46 |
|  | Robin | 1099 | 70 | 734 | 56 | 639 | 52 | 1120 | 68 | 1491 | 79 |
| Time of the year | Pre-breeding | 1698 | 128(12) | 1300 | 106(9) | 944 | 85(7) | 1754 | 131(11) | 2528 | 152(12) |
|  | Post-breeding | 2157 | 134(12) | 1301 | 107(9) | 1024 | 84(7) | 2031 | 143(11) | 2578 | 148(12) |
| Year | 2020 | 2157 | 136(9) | 1448 | 107(6) | 1146 | 84(4) | 2140 | 142(8) | 2784 | 151(9) |
|  | 2021 | 1698 | 126(9) | 1153 | 106(6) | 822 | 85(4) | 1645 | 132(8) | 2322 | 149(9) |

**Table S2**. Comparison of two modelling approaches for determining activity onset. Model coefficient values (‘Estimate’) and 95% Credible Intervals (95%CrI) for predictors explaining variation in relative time of onset of activity (i.e., time of onset of activity minus sunrise time; n = 3,855 observations of daily onset of activity for the frequentist approach; n = 3,476 observations of daily onset of activity for the Bayesian approach; note that the sample sizes differ slightly due to differences in accurate estimations provided by each method). Two models were fitted using, as the response variable, estimates of onset of activity extracted via a frequentist change point analysis or a Bayesian broken stick model (see Supplementary methods 1 and Supplementary results 1). Both methods produced in general consistent results (see also Figure 1b and Figure S11a). Model coefficients for relative onset of activity are given in hours after sunrise (posterior median given for estimates of Bayesian approach). Estimates for random effect intercepts are standard deviations. Reference level (i.e., intercept) is for post-breeding forest blackbirds in 2020.

| **Fixed effects** | **Estimate** | | **95%CrI** | |
| --- | --- | --- | --- | --- |
|  | **Frequentist approach** | **Bayesian approach** | **Frequentist approach** | **Bayesian approach** |
| Intercept | 50.577 | 75.388 | -138.499, 238.912 | -126.305, 283.196 |
| Time of the year |  |  |  |  |
| *Pre-breeding period* | -0.837 | -0.845 | -0.962, -0.714 | -0.981, -0.710 |
| Habitat |  |  |  |  |
| Urban | -0.534 | -0.561 | -0.978, -0.136 | -1.105, -0.059 |
| Species |  |  |  |  |
| *Blue tit* | 0.278 | 0.164 | -0.035, 0.600 | -0.221, 0.541 |
| *Chaffinch* | 0.221 | -0.129 | -0.123, 0.570 | -0.520, 0.291 |
| *Dunnock* | 0.152 | 0.120 | -0.164, 0.476 | -0.275, 0.513 |
| *Great tit* | 0.217 | 0.002 | -0.120, 0.563 | -0.390, 0.388 |
| *Robin* | -0.256 | -0.317 | -0.563, 0.054 | -0.680, 0.069 |
| Year | -0.025 | -0.037 | -0.118, 0.069 | -0.140, 0.063 |
| Minimum daily temperature | 0.004 | -0.001 | -0.005, 0.013 | -0.010, 0.008 |
| Daily rainfall | 0.000 | 0.001 | -0.003, 0.003 | -0.002, 0.004 |
| Habitat x Species |  |  |  |  |
| *Urban x Blue tit* | 0.273 | 0.334 | -0.139, 0.664 | -0.123, 0.793 |
| *Urban x Chaffinch* | 0.397 | 0.891 | -0.064, 0.855 | 0.383, 1.388 |
| *Urban x Dunnock* | 0.334 | 0.501 | -0.077, 0.740 | 0.046, 0.969 |
| *Urban x Great tit* | 0.555 | 0.728 | 0.139, 0.974 | 0.262, 1.191 |
| *Urban x Robin* | 0.044 | 0.187 | -0.337, 0.421 | -0.251, 0.620 |
| Habitat x Time of the year |  |  |  |  |
| *Pre-breeding period x Urban* | -0.028 | -0.089 | -0.205, 0.167 | -0.266, 0.096 |
| **Random effects and**  **residual terms** | **Estimate** | | **95%CrI** | |
|  | **Frequentist approach** | **Bayesian approach** | **Frequentist approach** | **Bayesian approach** |
| Randon intercept: Bird ID Forest | 0.271 | 0.350 | 0.221, 0.328 | 0.285, 0.426 |
| Randon intercept: Bird ID Urban | 0.373 | 0.446 | 0.313, 0.442 | 0.386, 0.514 |
| Randon intercept: Site | 0.096 | 0.142 | 0.002, 0.474 | 0.005, 0.530 |
| Randon intercept: Date | 0.061 | 0.095 | 0.012, 0.100 | 0.062, 0.130 |
| Residual standard deviation: Forest | -0.539 | -0.621 | -0.576, -0.503 | -0.660, -0.581 |
| Residual standard deviation: Urban | -0.172 | -0.390 | -0.201, -0.141 | -0.423, -0.358 |

**Table S3**. Comparison of two modelling approaches for determining activity end. Model coefficient values (‘Estimate’) and 95% Credible Intervals (95%CrI) for predictors explaining variation in relative time of end of activity (i.e., time of end of activity minus sunset time; n = 2,601 observations of daily end of activity for the frequentist approach; n = 3,198 observations of daily end of activity for the Bayesian approach). Two models were fitted using, as the response variable, estimates of end of activity extracted via a frequentist change point analysis or a Bayesian broken stick model (see Supplementary methods 1 and Supplementary results 1). Both methods produced in general consistent results (see also Figure 1d and Figure S11b). Model coefficients for relative end of activity are given in hours after sunset (posterior median given for estimates of Bayesian approach). Estimates for random effect intercepts are standard deviations. Reference level (i.e., intercept) is for post-breeding forest blackbirds in 2020.

| **Fixed effects** | **Estimate** | | **95%CrI** | |
| --- | --- | --- | --- | --- |
|  | **Frequentist approach** | **Bayesian approach** | **Frequentist approach** | **Bayesian approach** |
| Intercept | -53.928 | -30.028 | -283.669, 175.098 | -226.401, 163.919 |
| Time of the year |  |  |  |  |
| *Pre-breeding period* | -1.034 | -1.123 | -1.181, -0.882 | -1.279, -0.967 |
| Habitat |  |  |  |  |
| Urban | 0.392 | 0.107 | -0.077, 0.866 | -0.531, 0.703 |
| Species |  |  |  |  |
| *Blue tit* | -0.112 | -0.339 | -0.439, 0.229 | -0.805, 0.100 |
| *Chaffinch* | -0.635 | -0.627 | -0.994, -0.269 | -1.090, -0.170 |
| *Dunnock* | 0.023 | -0.062 | -0.316, 0.367 | -0.520, 0.403 |
| *Great tit* | -0.255 | -0.301 | -0.615, 0.109 | -0.748, 0.137 |
| *Robin* | 0.356 | 0.295 | 0.024, 0.692 | -0.146, 0.716 |
| Year | 0.027 | 0.016 | -0.086, 0.141 | -0.080, 0.113 |
| Minimum daily temperature | -0.024 | -0.006 | -0.037, -0.010 | -0.017, 0.004 |
| Daily rainfall | -0.002 | 0.000 | -0.008, 0.003 | -0.004, 0.004 |
| Habitat x Species |  |  |  |  |
| *Urban x Blue tit* | -0.287 | -0.049 | -0.722, 0.114 | -0.538, 0.442 |
| *Urban x Chaffinch* | 0.150 | 0.099 | -0.344, 0.654 | -0.409, 0.603 |
| *Urban x Dunnock* | -0.254 | -0.323 | -0.666, 0.165 | -0.820, 0.161 |
| *Urban x Great tit* | -0.359 | -0.327 | -0.775, 0.080 | -0.797, 0.151 |
| *Urban x Robin* | -0.176 | -0.149 | -0.589, 0.221 | -0.602, 0.314 |
| Habitat x Time of the year |  |  |  |  |
| *Pre-breeding period x Urban* | -0.100 | 0.225 | -0.293, 0.090 | 0.045, 0.405 |
| **Random effects and**  **residual terms** | **Estimate** | | **95%CrI** | |
|  | **Frequentist approach** | **Bayesian approach** | **Frequentist approach** | **Bayesian approach** |
| Randon intercept: Bird ID Forest | 0.192 | 0.373 | 0.120, 0.275 | 0.276, 0.484 |
| Randon intercept: Bird ID Urban | 0.291 | 0.255 | 0.229, 0.362 | 0.198, 0.318 |
| Randon intercept: Site | 0.119 | 0.161 | 0.003, 0.540 | 0.005, 0.689 |
| Randon intercept: Date | 0.153 | 0.105 | 0.107, 0.200 | 0.063, 0.145 |
| Residual standard deviation: Forest | -0.326 | -0.401 | -0.369, -0.283 | -0.444, -0.356 |
| Residual standard deviation: Urban | -0.258 | -0.324 | -0.297, -0.219 | -0.358, -0.289 |

**Table S4**. Analysis of activity duration. Model coefficient values (‘Estimate’) and 95% Credible Intervals (95%CrI) for predictors explaining variation in duration of diurnal activity (i.e., time between onset and end of daily activity; n = 1,968 observations of duration of diurnal activity). Model coefficients are given in hours. Estimates for random effect intercepts are standard deviations. Reference level (i.e., intercept) is for post-breeding forest blackbirds in 2020.

| **Fixed effects** | **Estimate**  **(posterior median)** | **95%CrI** |
| --- | --- | --- |
| Intercept | 2,109.664 | 1,488.197, 2,736.896 |
| Time of the year |  |  |
| *Pre-breeding period* | 1.222 | 0.847, 1.631 |
| Habitat |  |  |
| Urban | 0.366 | -0.359, 1.059 |
| Species |  |  |
| *Blue tit* | -0.547 | -0.937, -0.169 |
| *Chaffinch* | 0.122 | -0.203, 0.423 |
| *Dunnock* | 0.176 | -0.160, 0.553 |
| *Great tit* | 0.366 | -0.359, 1.059 |
| *Robin* | 1.067 | 0.790, 1.328 |
| Year | -1.039 | -1.349, -0.731 |
| Minimum daily temperature | 0.067 | 0.024, 0.112 |
| Daily rainfall | -0.008 | -0.019, 0.004 |
| Habitat x Species | 0.302 | -0.481, 1.112 |
| *Urban x Blue tit* | -0.038 | -0.643, 0.570 |
| *Urban x Chaffinch* | -0.634 | -1.253, -0.030 |
| *Urban x Dunnock* | 0.357 | -0.184, 0.888 |
| *Urban x Great tit* | 0.302 | -0.481, 1.112 |
| *Urban x Robin* | -0.038 | -0.643, 0.570 |
| Habitat x Time of the year |  |  |
| *Pre-breeding period x Urban* | -0.258 | -0.655, 0.115 |
| **Random effects and**  **residual terms** | **Estimate**  **(posterior median)** | **95%CrI** |
| Randon intercept: Bird ID Forest | 0.327 | 0.195, 0.485 |
| Randon intercept: Bird ID Urban | 0.605 | 0.480, 0.759 |
| Randon intercept: Site | 0.257 | 0.008, 1.081 |
| Randon intercept: Date | 0.769 | 0.647, 0.911 |
| Random intercept: Observation ID Forest | -0.100 | -0.152, -0.048 |
| Random intercept: Observation ID Urban | 0.160 | 0.115, 0.206 |

**Table S5**. Analysis of daytime activity level. Model coefficient values (‘Estimate’) and 95% Credible Intervals (95%CrI) for predictors explaining variation in levels of diurnal activity (i.e., proportion of time active during the diurnal phase of the day; n = 3,785 individual-days of observations). Model coefficients are given in the link scale (logit) after fitting a generalised linear mixed model with binomial error structure. An observation-level random effect was included to quantify within individual variation. Estimates for random effect intercepts are standard deviations. Reference level (i.e., intercept) is for post-breeding forest blackbirds in 2020.

| **Fixed effects** | **Estimate**  **(posterior median)** | **95%CrI** |
| --- | --- | --- |
| Intercept | 71.187 | -120.899, 263.748 |
| Time of the year |  |  |
| *Pre-breeding period* | -0.077 | -0.232, 0.080 |
| Habitat |  |  |
| Urban | -0.084 | -0.655, 0.836 |
| Species |  |  |
| *Blue tit* | 0.677 | 0.012, 1.342 |
| *Chaffinch* | 0.364 | -0.329, 1.077 |
| *Dunnock* | 0.184 | -0.517, 0.882 |
| *Great tit* | 0.565 | -0.129, 1.240 |
| *Robin* | 0.592 | -0.058, 1.238 |
| Year | -0.035 | -0.131, 0.060 |
| Minimum daily temperature | -0.006 | -0.012, 0.000 |
| Daily rainfall | -0.002 | -0.005, 0.000 |
| Habitat x Species |  |  |
| *Urban x Blue tit* | -0.227 | -0.928, 0.476 |
| *Urban x Chaffinch* | -0.144 | -0.962, 0.632 |
| *Urban x Dunnock* | -0.113 | -0.872, 0.641 |
| *Urban x Great tit* | -0.125 | -0.827, 0.602 |
| *Urban x Robin* | -0.279 | -0.963, 0.406 |
| Habitat x Time of the year |  |  |
| *Pre-breeding period x Urban* | -0.002 | -0.188, 0.181 |
| **Random effects and**  **residual terms** | **Estimate**  **(posterior median)** | **95%CrI** |
| Randon intercept: Bird ID Forest | 0.661 | 0.568, 0.770 |
| Randon intercept: Bird ID Urban | 0.478 | 0.404, 0.560 |
| Randon intercept: Site | 0.148 | 0.004, 0.619 |
| Randon intercept: Date | 0.061 | 0.038, 0.084 |
| Random intercept: Observation ID Forest | 0.509 | 0.486, 0.534 |
| Random intercept: Observation ID Urban | 0.272 | 0.257, 0.288 |

**Table S6**. Analysis of nighttime activity level. Model coefficient values (‘Estimate’) and 95% Credible Intervals (95%CrI) for predictors explaining variation in levels of nocturnal activity (i.e., proportion of time active during the nocturnal phase of the day; n = 5,106 individual-days of observations). Model coefficients are given in the link scale (logit) after fitting a zero-inflated generalised linear mixed model with binomial error structure. An observation-level random effect was included to quantify within individual variation. Estimates for random effect intercepts are standard deviations. Reference level (i.e., intercept) is for post-breeding forest blackbirds in 2020.

| **Fixed effects** | **Estimate**  **(posterior median)** | **95%CrI** |
| --- | --- | --- |
| Zero-inflation intercept | -1.888 | -2.275, -1.570 |
| Intercept | -328.872 | -994.080, 303.807 |
| Time of the year |  |  |
| *Pre-breeding period* | -0.434 | -0.991, 0.140 |
| Habitat |  |  |
| Urban | 2.684 | 0.832, 4.561 |
| Species |  |  |
| *Blue tit* | 0.776 | -0.757, 2.357 |
| *Chaffinch* | 1.093 | -0.483, 2.736 |
| *Dunnock* | -0.610 | -2.272, 1.003 |
| *Great tit* | 1.102 | -0.527, 2.757 |
| *Robin* | 0.214 | -1.294, 1.716 |
| Year | 0.160 | -0.153, 0.489 |
| Minimum daily temperature | -0.013 | -0.046, 0.019 |
| Daily rainfall | 0.006 | -0.007, 0.018 |
| Habitat x Species |  |  |
| *Urban x Blue tit* | -0.827 | -2.501, 0.860 |
| *Urban x Chaffinch* | -1.763 | -3.596, 0.025 |
| *Urban x Dunnock* | 0.400 | -1.345, 2.127 |
| *Urban x Great tit* | -1.726 | -3.477, 0.045 |
| *Urban x Robin* | -0.369 | -1.927, 1.283 |
| Habitat x Time of the year |  |  |
| *Pre-breeding period x Urban* | -1.005 | -1.611, -0.401 |
| **Random effects and**  **residual terms** | **Estimate**  **(posterior median)** | **95%CrI** |
| Randon intercept: Bird ID Forest | 1.427 | 1.166, 1.729 |
| Randon intercept: Bird ID Urban | 1.159 | 1.009, 1.331 |
| Randon intercept: Site | 0.471 | 0.013, 1.699 |
| Randon intercept: Date | 0.414 | 0.330, 0.508 |
| Random intercept: Observation ID Forest | 1.427 | 1.445, 1.757 |
| Random intercept: Observation ID Urban | 1.084 | 1.008, 1.165 |

**Table S7**. Bivariate analysis of diurnal and nocturnal activity. Model coefficient values (‘Estimate’) and 95% Credible Intervals (95%CrI) for predictors explaining variation in levels of diurnal and nocturnal activity (i.e., proportion of time active during the diurnal and nocturnal phase of the day; n = 5,871 individual-days of observations) in a bivariate model. This model was fitted to estimate between- and within-individual correlations in diurnal and nocturnal activity level. Model coefficients are given in the link scale (logit) after fitting a bivariate generalised linear mixed model with binomial error structure. An observation-level random effect was included to quantify within individual variation. Estimates for random effect intercepts are standard deviations. Reference level (i.e., intercept) is for post-breeding forest blackbirds in 2020.

|  | **Diurnal activity** | | **Nocturnal activity** | |
| --- | --- | --- | --- | --- |
| **Fixed effects** | **Estimate** | **95%CrI** | **Estimate** | **95%CrI** |
| Zero-inflation intercept |  |  | -1.952 | -2.396, -1.573 |
| Intercept | 31.538 | -153.747, 203.748 | -360.943 | -1141.972, 414.888 |
| Time of the year |  |  |  |  |
| *Pre-breeding period* | -0.173 | -0.313, -0.038 | -0.526 | -1.330, 0.128 |
| Habitat |  |  |  |  |
| Urban | 0.167 | -0.560, 0.896 | 2.191 | -1.777, 4.472 |
| Species |  |  |  |  |
| *Blue tit* | 0.630 | -0.048, 1.266 | 0.637 | -0.951, 2.386 |
| *Chaffinch* | 0.181 | -0.457, 0.836 | 1.374 | -0.488, 3.259 |
| *Dunnock* | 0.233 | -0.462, 0.862 | -0.800 | -2.438, 0.998 |
| *Great tit* | 0.523 | 0.339, -0.143 | 0.837 | -0.998, 2.764 |
| *Robin* | 0.643 | -0.078, 1.262 | 0.073 | -1.505, 1.745 |
| Year | -0.016 | -0.101, 0.076 | 0.176 | -0.208, 0.562 |
| Minimum daily temperature | -0.008 | -0.013, -0.003 | -0.010 | -0.044, 0.023 |
| Daily rainfall | -0.002 | -0.005, -0.000 | 0.005 | -0.008, 0.019 |
| Habitat x Species |  |  |  |  |
| *Urban x Blue tit* | -0.155 | -0.791, 0.543 | -0.359 | -2.223, 1.489 |
| *Urban x Chaffinch* | -0.062 | -0.771, 0.629 | -0.437 | -2.696, 1.815 |
| *Urban x Dunnock* | -0.192 | -0.876, 0.439 | 0.822 | -1.123, 2.644 |
| *Urban x Great tit* | -0.012 | -0.688, 0.739 | -1.157 | -3.241, 0.781 |
| *Urban x Robin* | -0.420 | -1.087, 0.376 | 0.304 | -1.512, 2.052 |
| Habitat x Time of the year |  |  |  |  |
| *Pre-breeding period x Urban* | 0.086 | -0.074, 0.251 | -0.959 | -1.703, 0.059 |
| **Random effects** | **Estimate** | **95%CrI** | **Estimate** | **95%CrI** |
| Randon intercept: Bird ID Forest | 0.585 | 0.479, 0.709 | 1.384 | 1.085, 1.732 |
| Randon intercept: Bird ID Urban | 0.326 | 0.277, 0.387 | 1.241 | 1.042, 1.476 |
| Randon intercept: Site | 0.186 | 0.012, 0.725 | 0.550 | 0.017, 1.917 |
| Randon intercept: Date | 0.038 | 0.008, 0.063 | 0.391 | 0.289, 0.505 |
| Random intercept: Observation ID Forest | 0.355 | 0.334, 0.375 | 1.543 | 1.369, 1.733 |
| Random intercept: Observation ID Urban | 0.242 | 0.227, 0.258 | 0.987 | 0.902, 1.084 |
| **Between and within individual correlations** | **Estimate** | | **95%CrI** | |
| Correlation: Bird ID Forest | -0.233 | | -0.472, 0.030 | |
| Correlation: Bird ID Urban | 0.035 | | -0.192, 0.257 | |
| Correlation: Observation ID Forest | -0.152 | | -0.271, -0.043 | |
| Correlation: Observation ID Urban | -0.048 | | -0.150, 0.053 | |

**Supplementary Figures**

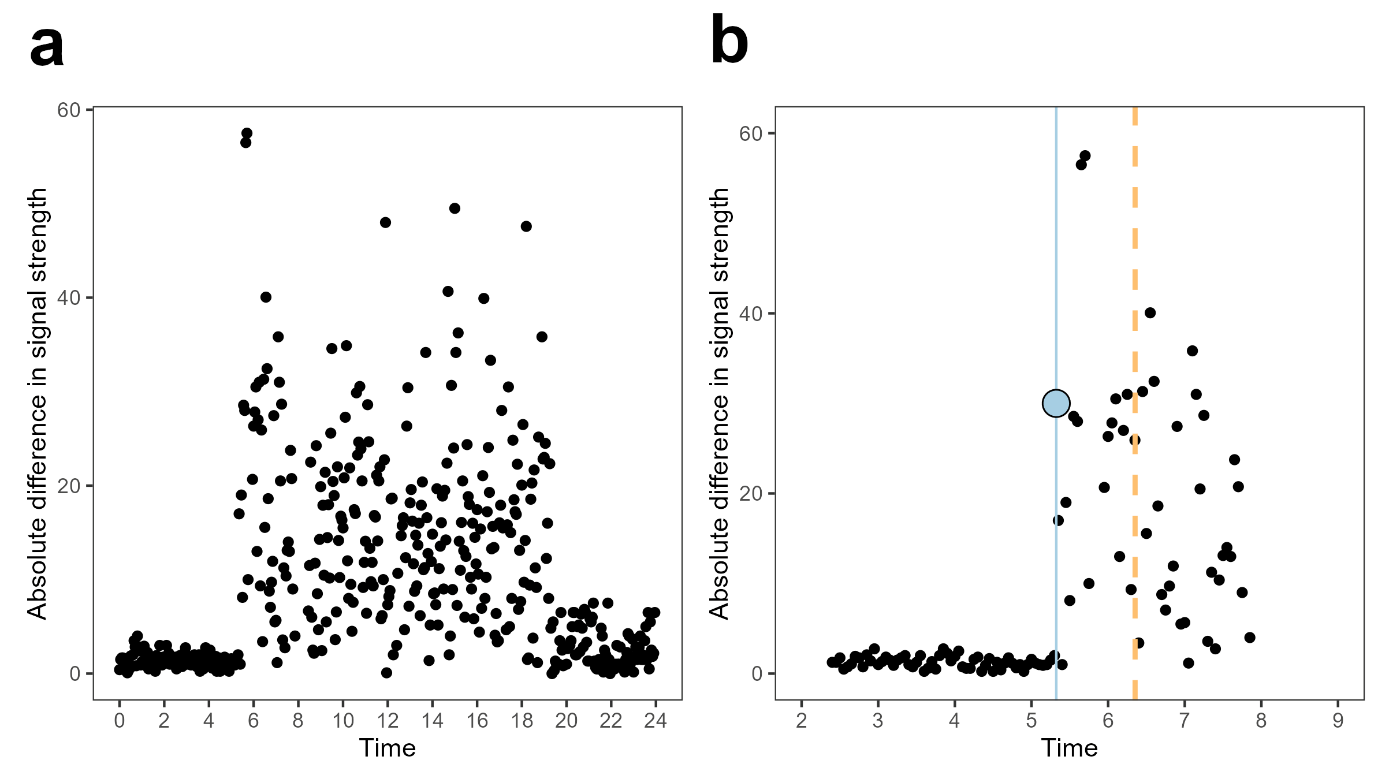

**Figure S1. One-day example of raw radio data collated and used to infer diel patterns of activity.** (**a**) Signal differentials (y-axis; absolute difference in signal strength between consecutive time points) were calculated throughout the day in 3-min bins. (**b**) Close-up of (a) around the time of activity onset. Changes in signal differential variation were used to infer time of onset of activity, here illustrated as the blue dot and vertical line (dashed yellow line illustrates sunrise time). A similar approach was employed to calculate the time of end of activity.

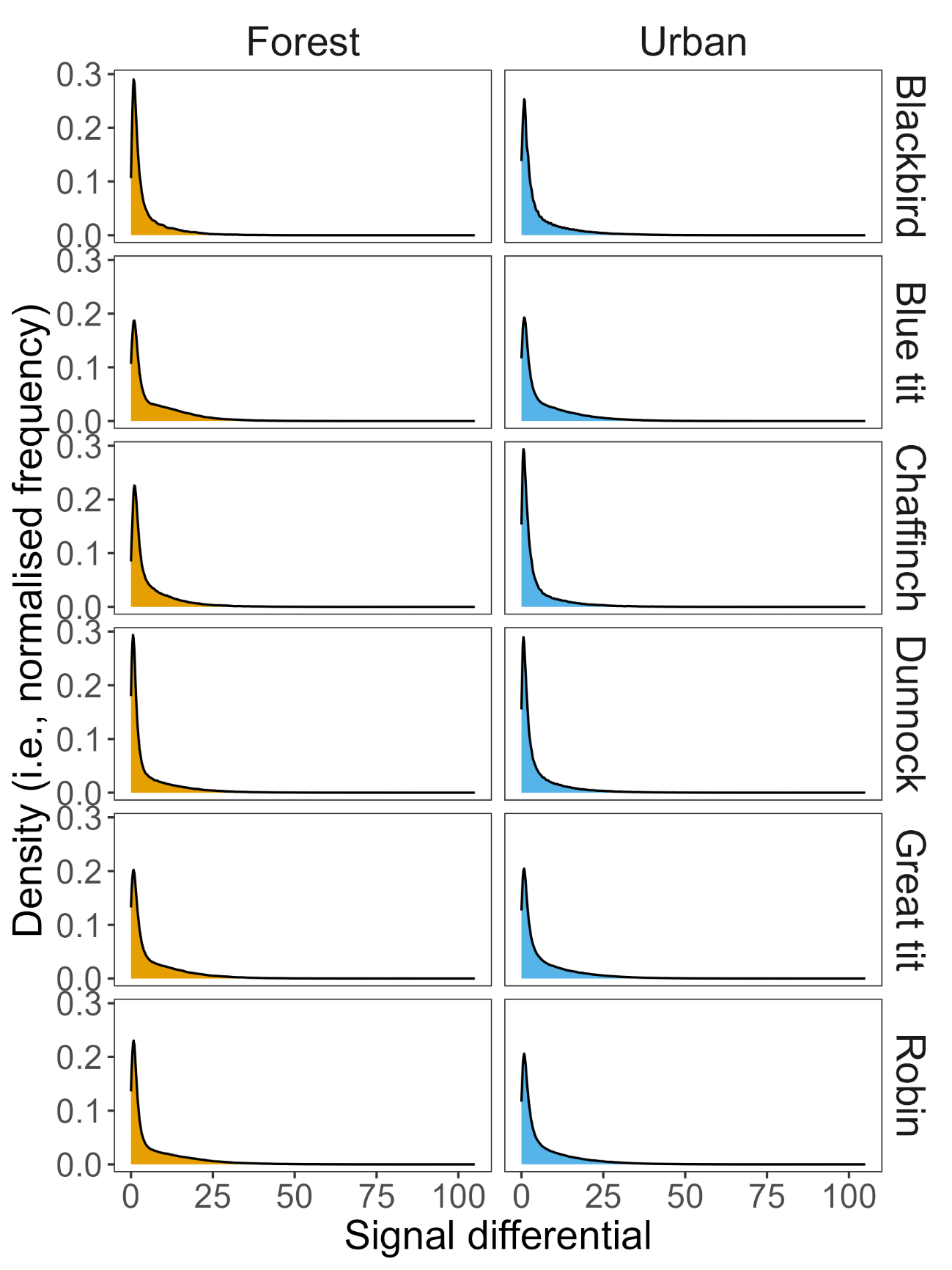

**Figure S2. Distribution of signal differentials across species and habitats.** The distribution of signal differentials showed similar patterns across species and habitats, with high frequency of low values and a low proportion of high values.

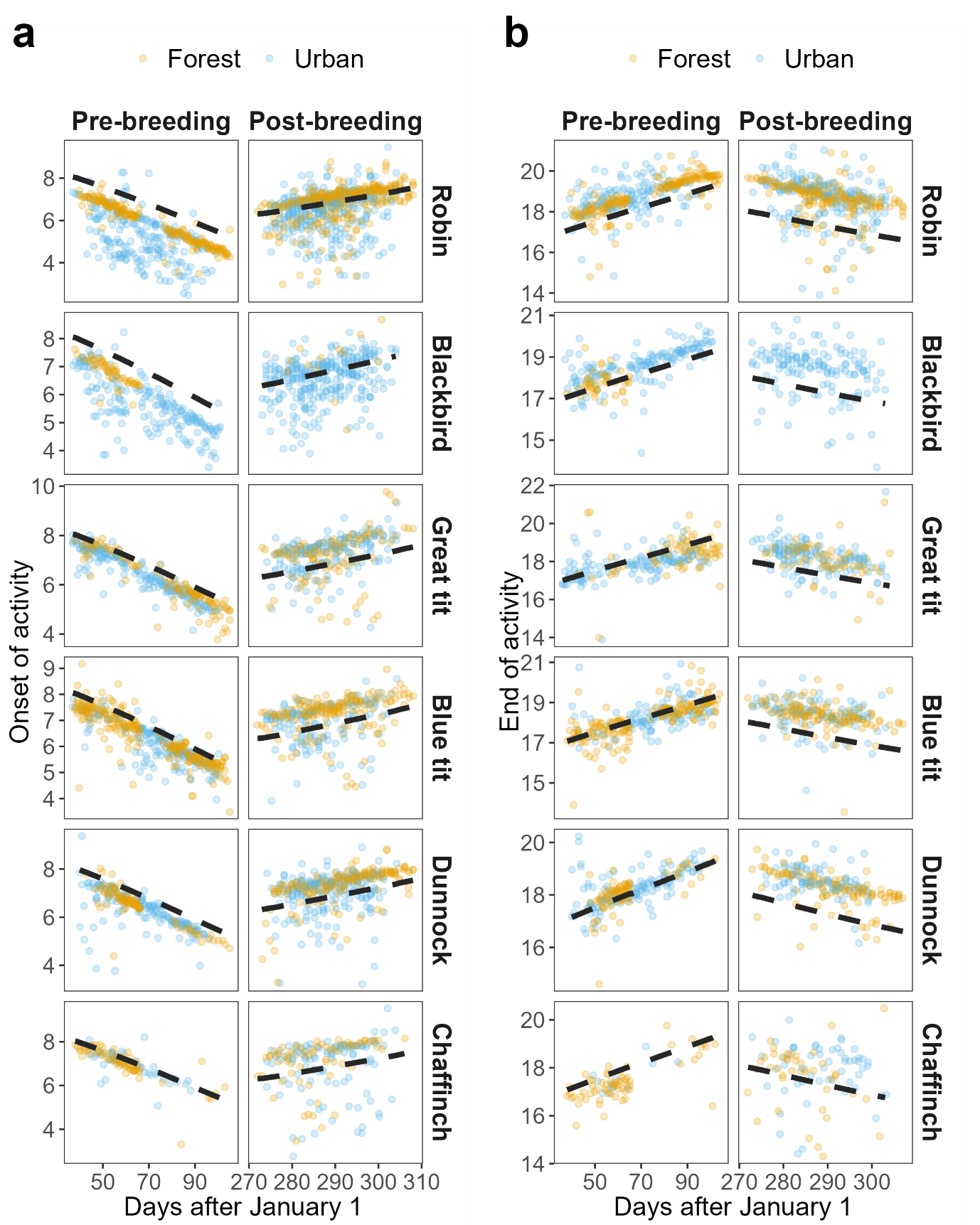

**Figure S3. Absolute time of onset and end of activity in urban and forest habitats throughout the two sampling periods across species during the pre-breeding phase (left) and post-breeding phase (right).** (**a**) absolute time of onset of activity. (**b**) Absolute time of end of activity. Dashed black line illustrates time of sunrise (in a) and time of sunset (in b).

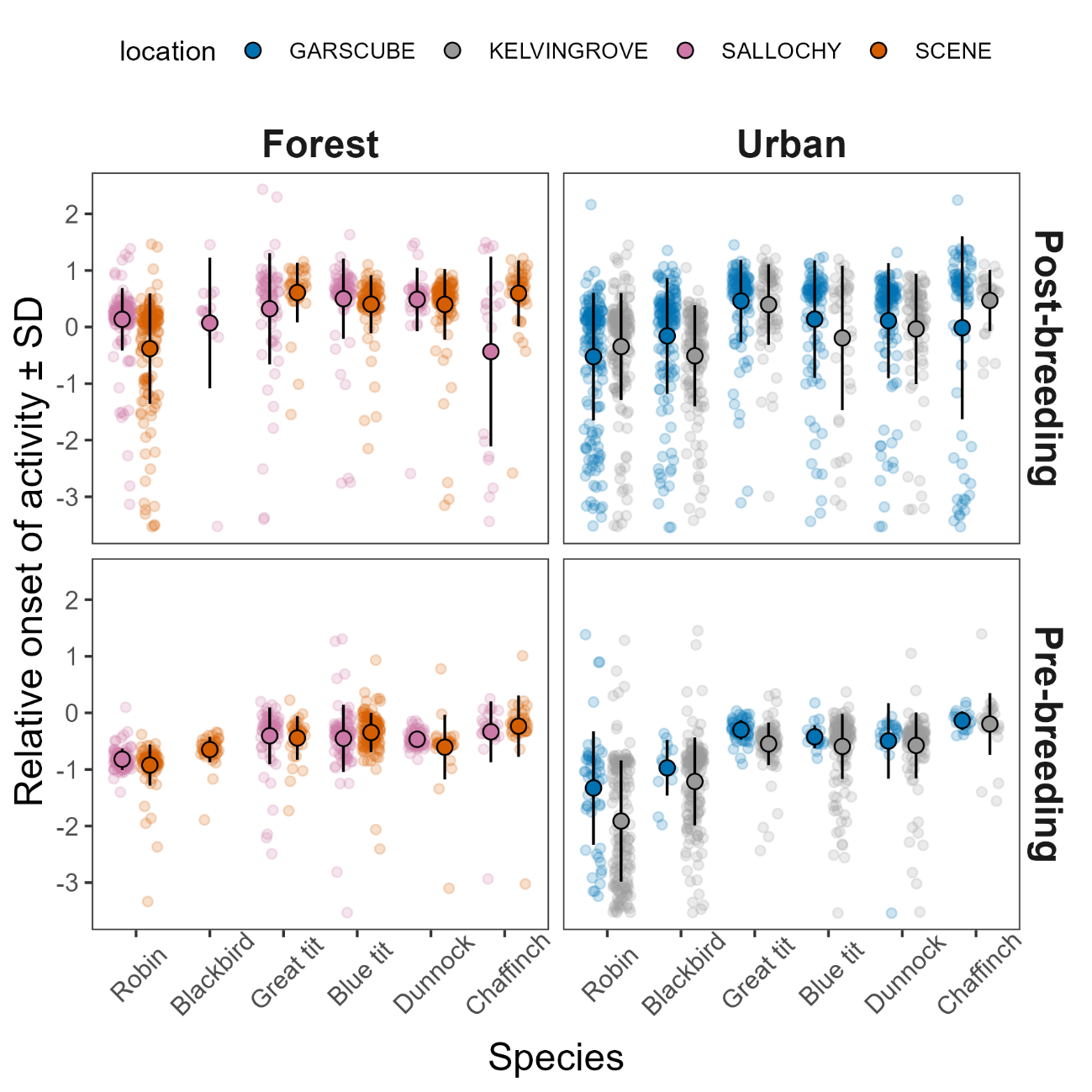

**Figure S4. Variation in relative onset of activity between sites within habitats.** Raw data points are illustrated by small translucid dots, while larger opaque points and whiskers provide raw mean values and standard deviations.

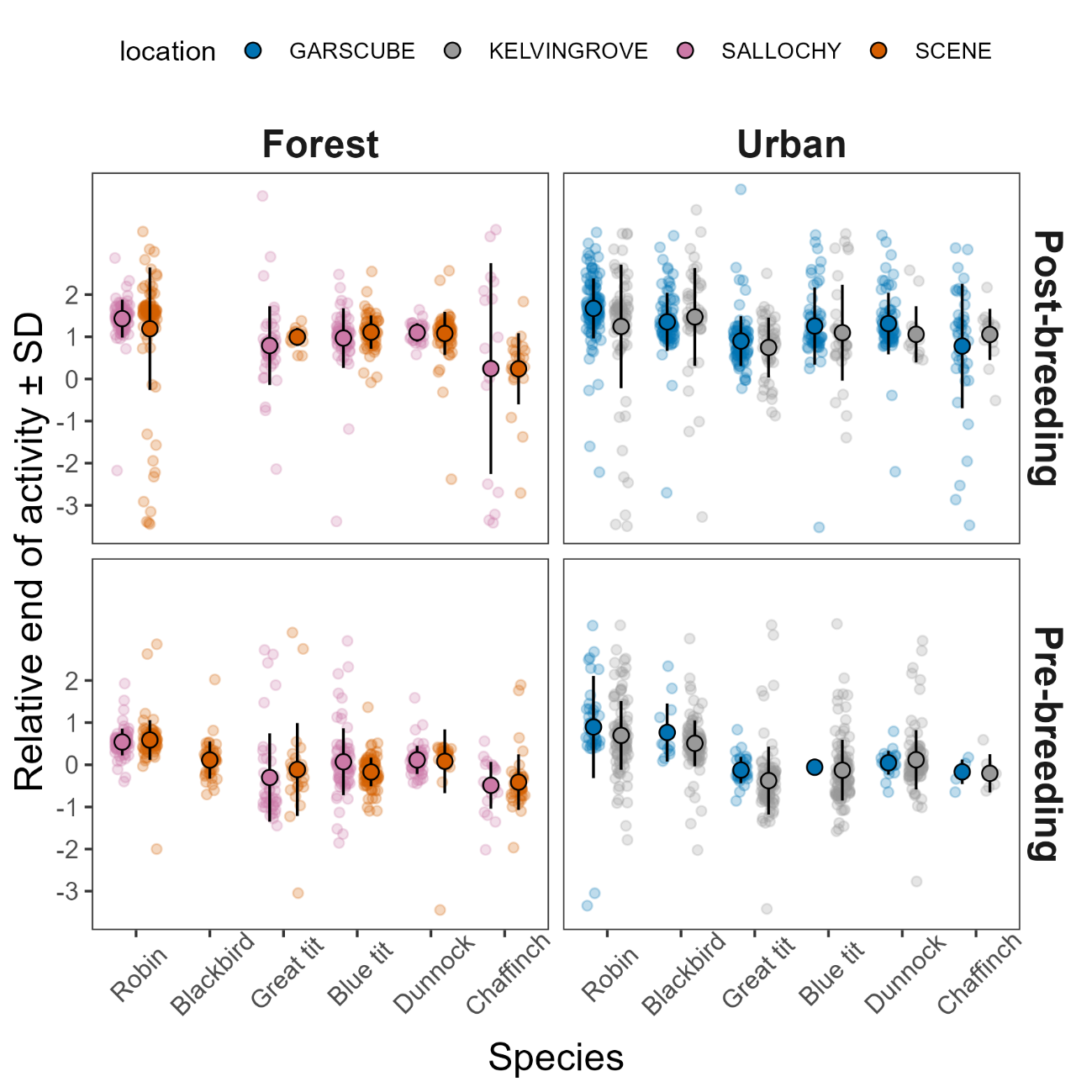

**Figure S5. Variation in relative end of activity between sites within habitats.** Raw data points are illustrated by small translucid dots, while larger opaque points and whiskers provide raw mean values and standard deviations.

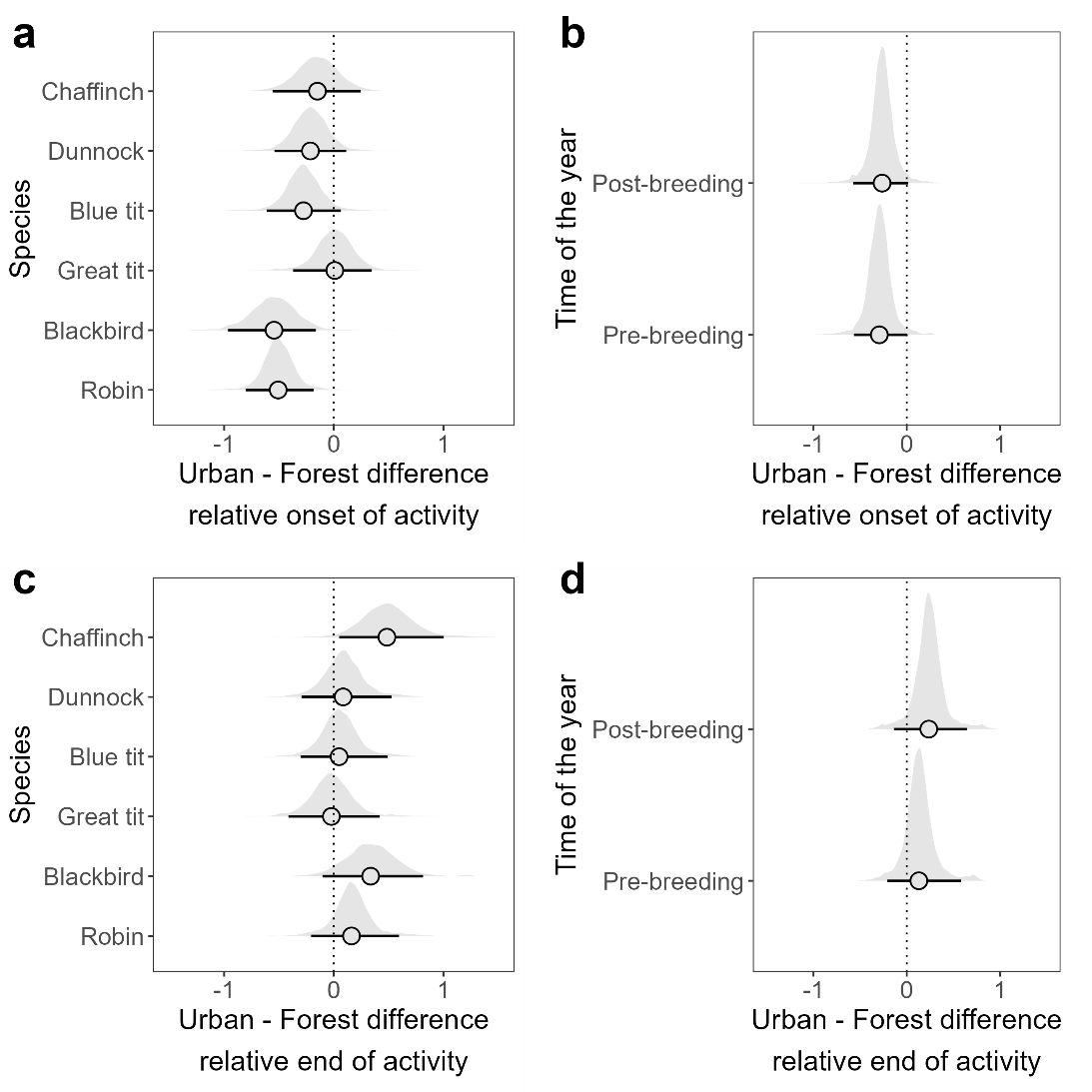

**Figure S6**. **Differences in onset and end of activity between urban and forest habitats across species and times of the year.** Dotted lines mark zero. (**a**) Difference in mean model predictions between urban and forest onset of activity across species while averaging across other fixed effects (i.e., expected marginal effect of habitat per species). Negative values indicate earlier onset of activity in urban habitats. Urban – forest differences were particularly large for robins (posterior difference estimate [95%CrI] = -0.507 [-0.788, -0.173]) and blackbirds (posterior difference estimate [95%CrI] = -0.545 [-0.970, -0.172]). (**b**) Difference in mean model predictions between urban and forest onset of activity across post- and pre-breeding periods while averaging over other fixed effects (i.e., expected marginal effect of habitat per time of the year). Urban – forest differences were similar across time periods. (**c**) Difference in mean model predictions between urban and forest end of activity across species while averaging across other fixed effects (i.e., expected marginal effect of habitat per species). Negative values indicate earlier end of activity in urban habitats compared to forest habitats. Urban – forest differences were particularly large for chaffinches (posterior difference estimate [95%CrI] = 0.485 [0.002, 0.946]). (**d**) Difference in mean model predictions between urban and forest end of activity across post- and pre-breeding periods while averaging over other fixed effects (i.e., expected marginal effect of habitat per time of the year). Urban – forest differences in end of activity showed credible interval that included zero in both the pre-breeding and post-breeding periods and were similar in both periods. Full posterior distributions are shown along with posterior medians and 95%CrIs.

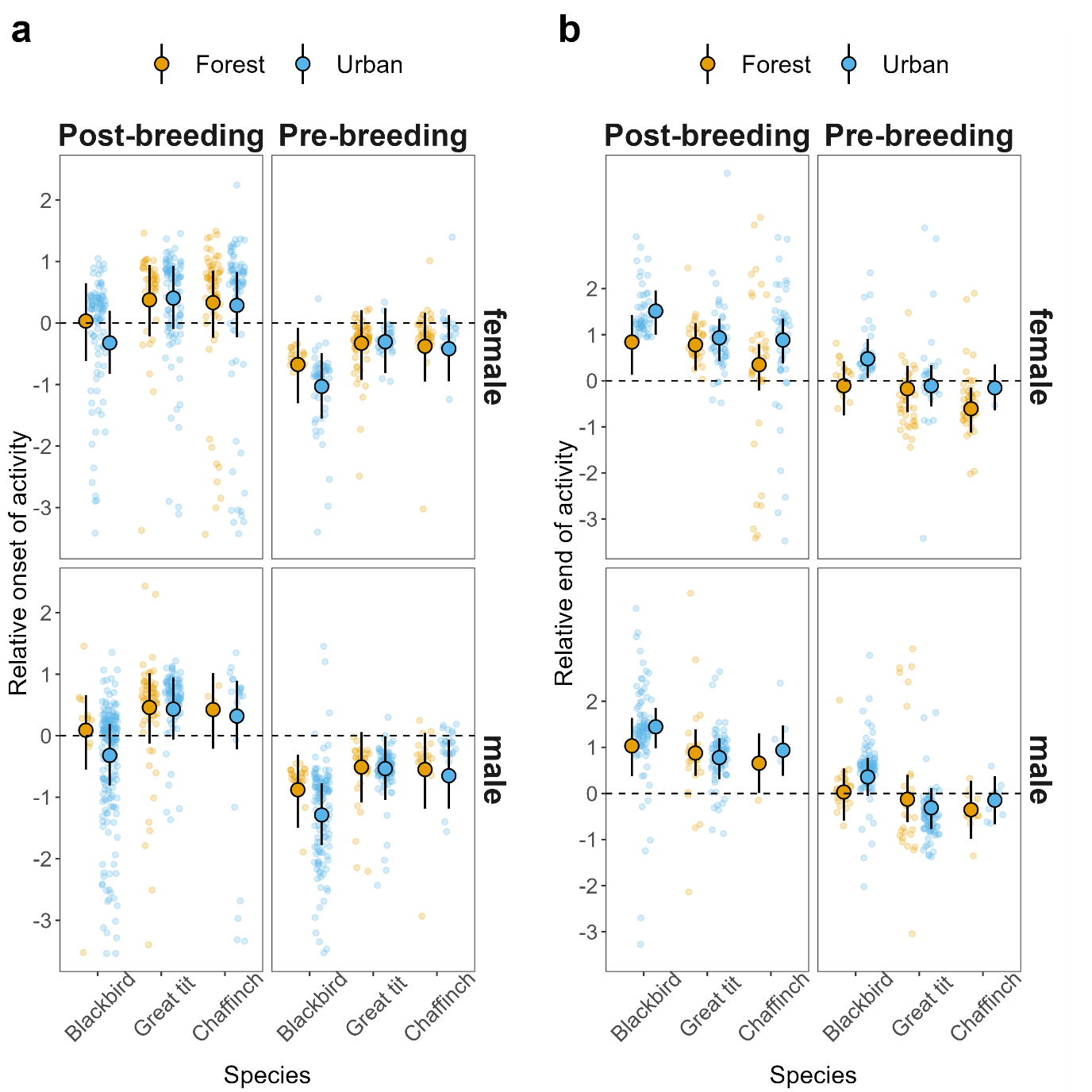

**Figure S7**. **Differences in onset and end of activity between urban and forest habitats across species, times of the year and sex.** (**a**) Relative onset and (**b**) relative end of activity in urban and forest habitats post- and pre-breeding across the three passerine species whose sex could be determined in the field from plumage characteristics. Urban effects on onset and end of activity were similar across both sexes. Raw data are shown as small translucid points with filled points and intervals illustrating expected model predictions and 95% Credible Interval (95%CrI).

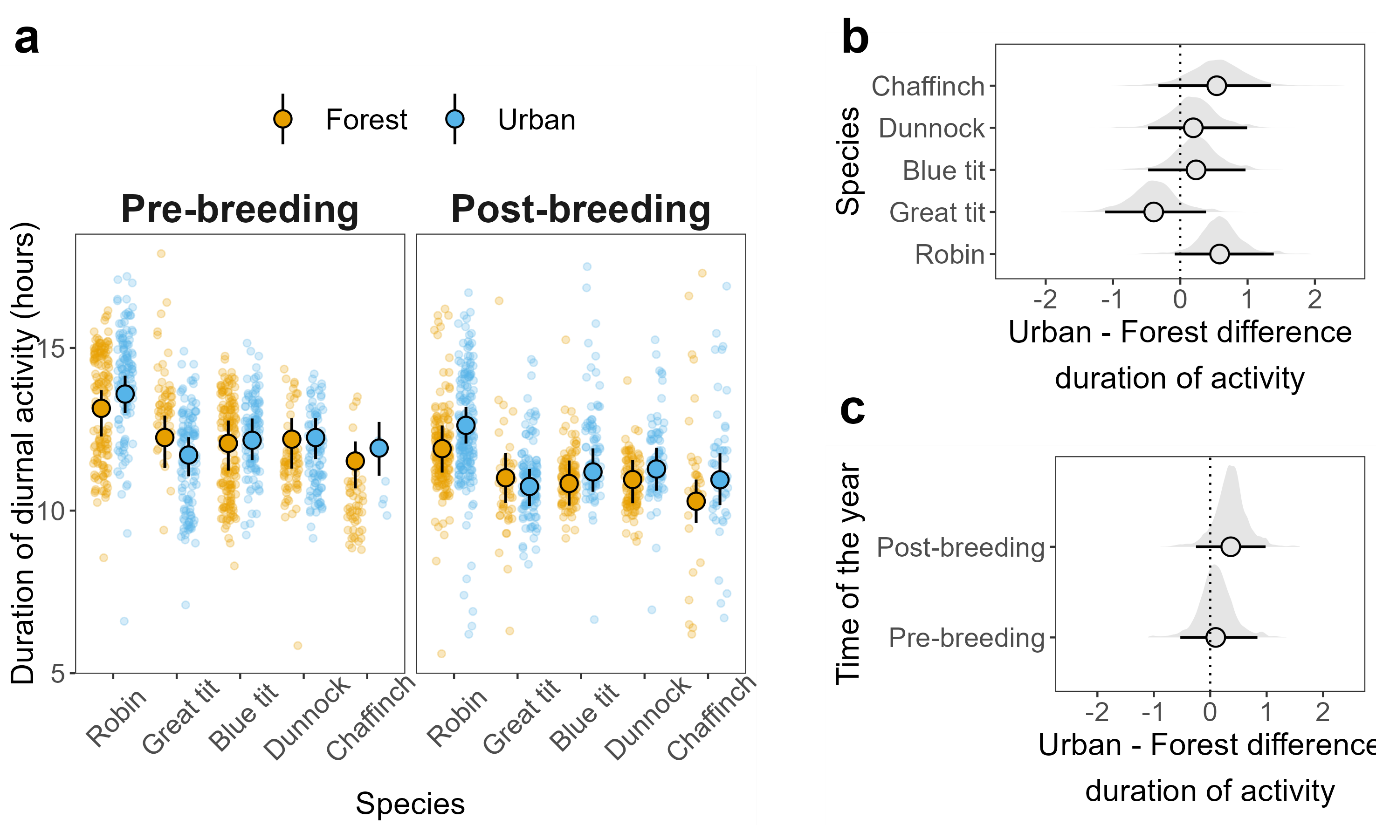

**Figure S8. Duration of diurnal activity in urban and forest habitats post- and pre-breeding across passerine species. (a)** Duration of activity in urban and forest habitats post- and pre-breeding across five passerine species (no data were available for blackbirds). Raw data are shown as small translucid points with filled points and intervals illustrating expected model predictions and 95% Credible Interval (95%CrI). (**b**) Difference in mean model predictions between urban and forest duration of activity across species while averaging across other fixed effects (i.e., expected marginal effect of habitat per species). Negative values indicate shorter durations in urban habitats, while positive values indicate longer durations in urban habitats compared to forest habitats. (**c**) Difference in mean model predictions between urban and forest duration of activity across post- and pre-breeding periods while averaging over other fixed effects (i.e., expected marginal effect of habitat per time of the year). The formal comparison of these effects yielded a non-significant difference (difference in urban effect pre-breeding minus post-breeding [95%CrI] = -0.254 [-0.655, +0.115]). Negative values indicate shorter durations in urban habitats, while positive values indicate longer durations in urban habitats compared to forest habitats. Dotted line in b and c mark zero (i.e., no difference in effect across habitats). Full posterior distributions are shown in b and c along with posterior medians and 95%CrIs.

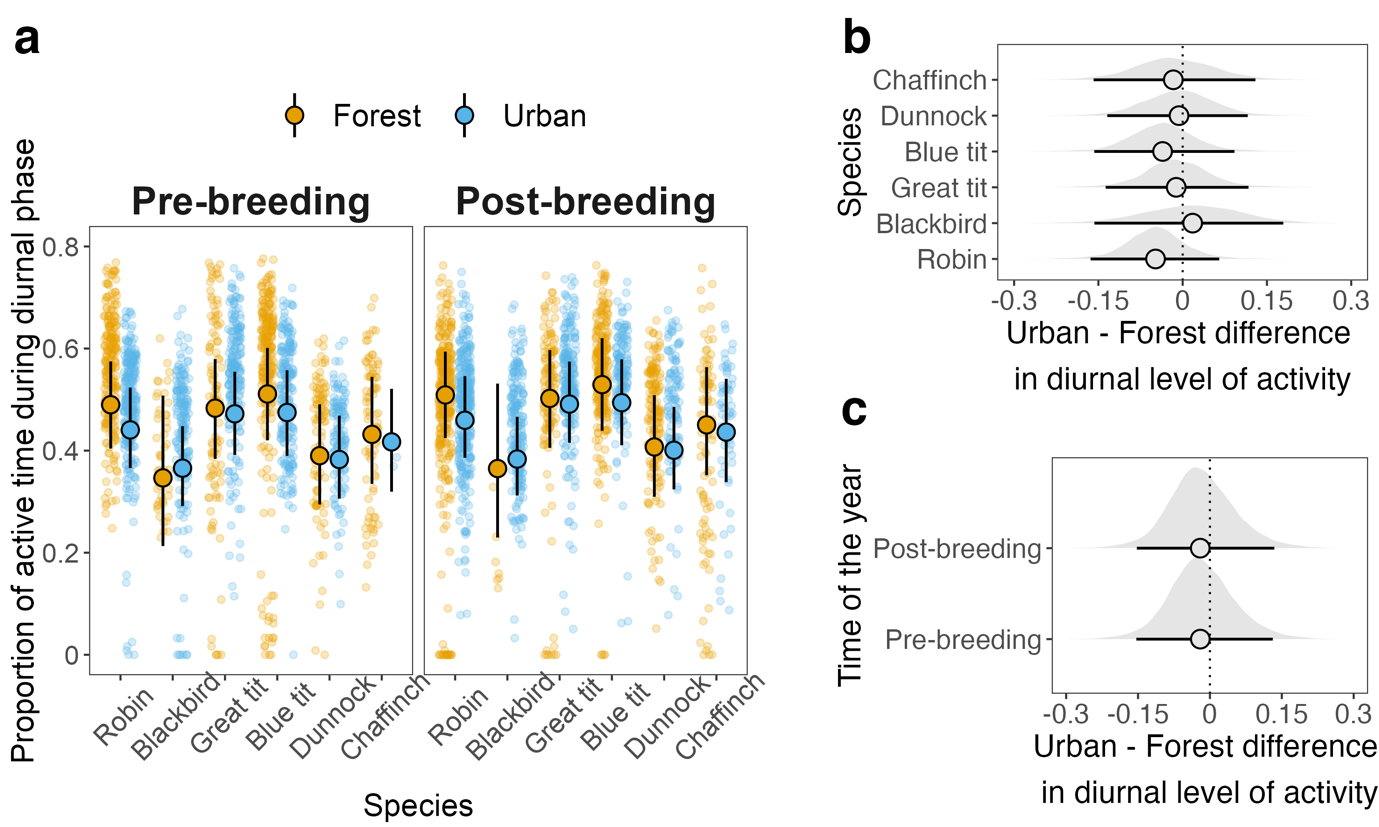

**Figure S9. Diurnal level of activity (proportion of time active during the diurnal phase) in urban and forest habitats post- and pre-breeding across passerine species. (a)** Diurnal level of activity in urban and forest habitats post- and pre-breeding across six passerine species. Raw data are shown as small translucid points with filled points and intervals illustrating expected model predictions and 95% Credible Interval (95%CrI). (**b**) Difference in expected mean model predictions between urban and forest diurnal level of activity across species while averaging across other fixed effects (i.e., expected marginal effect of habitat per species). Negative values indicate lower activity in urban habitats, while positive values indicate higher activity in urban habitats compared to forest habitats. (**c**) Difference in expected mean model predictions between urban and forest diurnal level of activity across post- and pre-breeding periods while averaging over other fixed effects (i.e., expected marginal effect of habitat per time of the year). Negative values indicate lower activity in urban habitats, while positive values indicate higher activity in urban habitats compared to forest habitats. Posterior distributions, medians and 95%CrIs for estimated proportions are presented in b and c. Dotted line in b and c marks zero (i.e., no difference in effect across habitats).

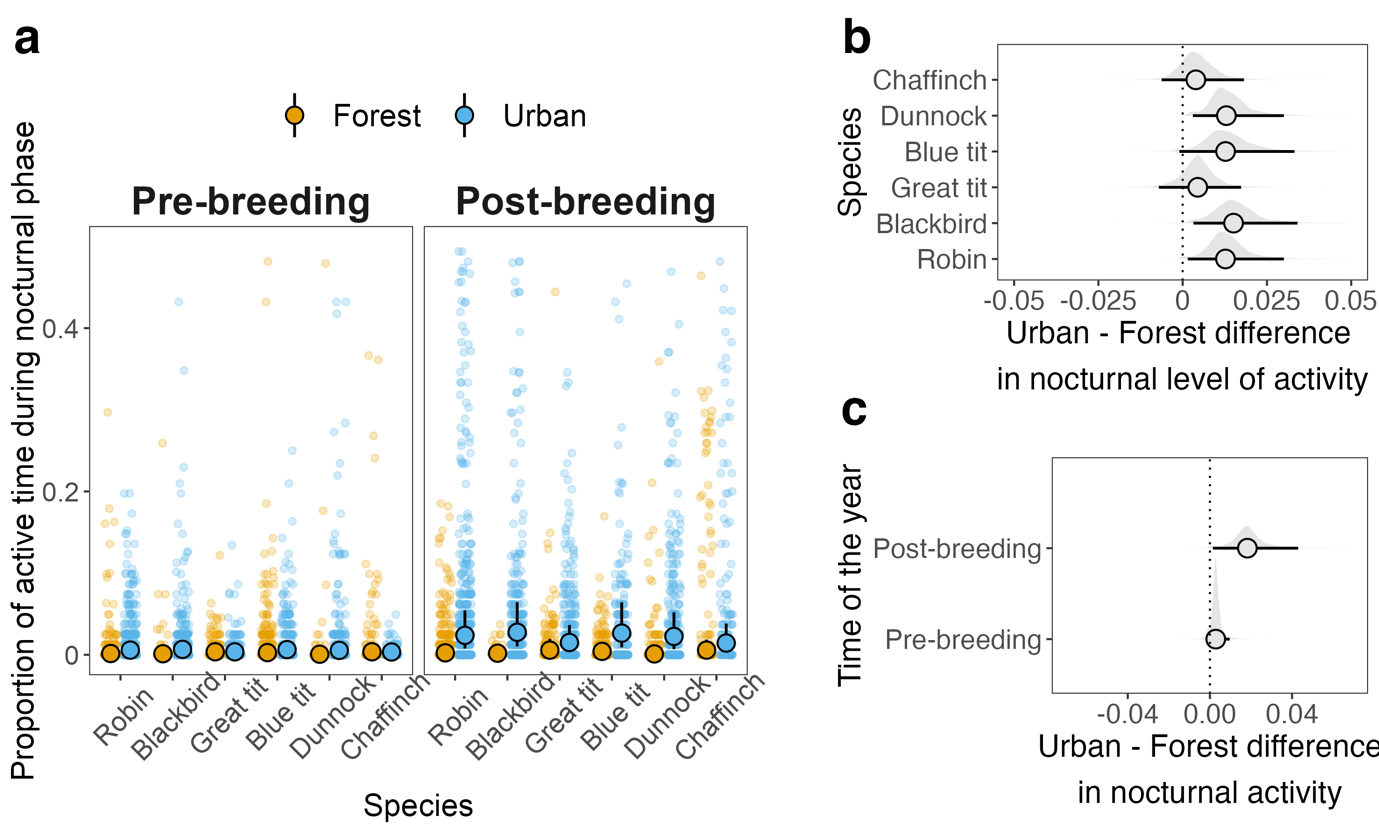

**Figure S10. Nocturnal level of activity (proportion of time active during the nocturnal phase) in urban and forest habitats post- and pre-breeding across passerine species**. (**a**) Nocturnal level of activity in urban and forest habitats post- and pre-breeding across six passerine species. Raw data are shown as small translucid points with filled points and intervals illustrating expected model predictions and 95% Credible Interval (95%CrI). (**b**) Difference in expected mean model predictions between urban and forest level of nocturnal activity across species while averaging across other fixed effects (i.e., expected marginal effect of habitat per species). Negative values indicate lower activity in urban habitats, while positive values indicate higher activity in urban habitats compared to forest habitats. (**c**) Difference in expected mean model predictions between urban and forest level of nocturnal activity across post- and pre-breeding periods while averaging over other fixed effects (i.e., expected marginal effect of habitat per time of the year). Negative values indicate lower activity in urban habitats, while positive values indicate higher activity in urban habitats compared to forest habitats. The urban increase in nocturnal activity was particularly strong in the post-breeding period (posterior estimate for the difference between urban and forest habitats [95%CrI] = +0.018 [+0.001, +0.045]); this increase was higher than in the pre-breeding period (difference in urban effect pre-breeding minus post-breeding [95%CrI] = -0.015 [-0.036, -0.003]). Posterior distributions, medians and 95%CrIs for estimated proportions are presented in b and c. Dotted line in b and c marks zero (i.e., no difference in effect across habitats).

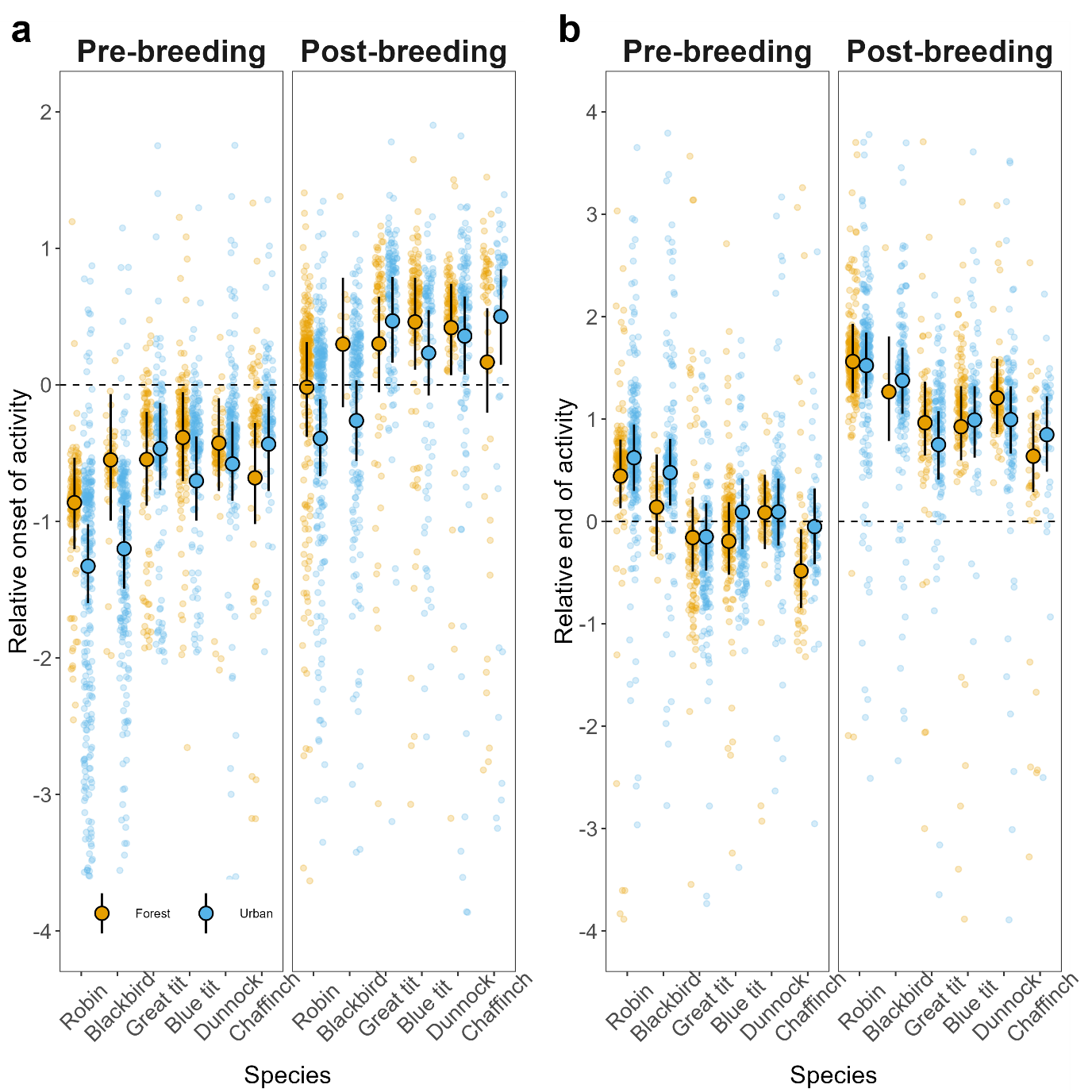

**Figure S11. Onset and end of activity in urban and forest habitats post- and pre-breeding across passerine species, after estimating time of onset and end of activity via Bayesian broken stick models (see Supplementary methods).** (**a**) Time of onset of activity relative to sunrise in urban and forest habitats pre- and post -breeding across the six species under study. (**b**) Time of end of activity relative to sunset in urban and forest habitats in pre- and post-breeding across the six species. These results, based on an alternative approach to estimate time of onset and end of activity, are in agreement with the results shown in the main text, where time of onset and end of activity were estimated using a frequentist analysis (Figure 1b and Figure 1d). Dashed lines in a and b represent sunrise time and sunset time, respectively. The raw data are shown as small translucid points with filled points and intervals illustrating model predictions and 95% Credible Interval (95%CrI). Blue shade always indicates urban populations, while yellow shade indicates forest populations.
